# The inner junction complex of the cilia is an interaction hub that involves tubulin post-translational modifications

**DOI:** 10.1101/774695

**Authors:** Ahmad Khalifa, Muneyoshi Ichikawa, Daniel Dai, Shintaroh Kubo, Corbin Black, Katya Peri, Thomas S. McAlear, Simon Veyron, Shun Kai Yang, Javier Vargas, Susanne Bechstedt, Jean-Francois Trempe, Khanh Huy Bui

## Abstract

Microtubules are cytoskeletal structures involved in structural support, microtubule-based transport and the organization of organelles in the cells. The building blocks of the microtubule, the α- and β-tubulin heterodimers, polymerize into protofilaments, that associate laterally to form the hollow microtubule. There exists a specific type of microtubule structures in the cilia, termed doublet microtubules, where high stability is required for ciliary beating and function. The doublet microtubule, consisting of a complete A-tubule and a partial B-tubule maintains its stability through unique interactions at its outer and inner junctions, where the A- and B-tubules meet.

Using cryo-electron microscopy, we present the answer to the long-standing question regarding the identities, localizations and structures of the *Chlamydomonas* doublet microtubule inner junction proteins. Using a combination of sequence bioinformatics and mass spectrometry, we identified two new inner junction proteins, FAP276 and FAP106, and an inner junction associated protein FAP126. We show that inner junction proteins PACRG and FAP20, together with FAP52, previously unidentified FAP276, FAP106 and FAP126, form an interaction hub at the inner junction, which involves tubulin sites for post-translational modifications. We further compare the *Chlamydomonas* and *Tetrahymena* doublet microtubule structures to understand the common and species-specific features of the inner junction.

## INTRODUCTION

Cilia and flagella are highly conserved organelles present in protists all the way to humans. They are commonly classified into two forms: motile and non-motile cilia. Motile cilia are responsible for mucus clearance in the airway, cerebrospinal fluid circulation and sperm motility (1). The non-motile cilia, namely primary cilia, function as the cellular antennas that sense chemical and mechanical changes. Cilia are essential for growth and development and therefore human health. Defects in cilia often result in abnormal motility or stability, which lead to cilia-related diseases such as primary ciliary dyskinesia, retinal degeneration, hydrocephalus and polydactyly (2).

Both cilia types are comprised of a bundle of nine specialized microtubule structures termed doublet microtubules (doublets). Ciliary components, important for motility such as the outer and inner dynein arms, radial spokes and the dynein regulatory complex (DRC) are assembled onto the surface of the doublet (3-6). Inside the doublets, is a weaving network of proteins, termed microtubule-inner-proteins (MIPs), that bind to the inner lumen surface of the doublet (7, 8). These MIPs act to stabilize the microtubule and very likely regulate the ciliary waveform through interactions with the tubulin lattice (8).

Doublets consists of a complete 13-protofilament (PF) A-tubule, similar to a 13-PF cytoplasmic microtubule and a partial 10-PF B-tubule forming on top of the A-tubule. To this day there still exists a long-standing question of how the junctions between the two tubules are formed (9-11). Recent high-resolution cryo-EM structure of the doublet shows that the outer junction is formed by a non-canonical tubulin interaction between PF B1 and PF A10 and A11 (7). The inner junction (IJ), which bridges the inner gap between the B-tubule and A-tubule is formed by non-tubulin proteins. Both primary and motile cilia have been observed to contain the IJ (11, 12).

In vitro formation of a B-tubule-like hook (i.e. the outer junction like interaction) was assembled onto pre-existing axonemal and mitotic spindle microtubule with the addition of purified brain tubulin (13). More recently, the B-tubule-like hook can be achieved by adding purified tubulins onto existing subtilisin treated microtubule (14). However, these hooks are not closed and appear to be very flexible (14). This supports the notion that the IJ is composed of non-tubulin proteins that are indispensable to the stability of the IJ.

The IJ is composed of FAP20 as shown through cryo-electron tomography and BCCP-tagging (15). Dymek et al (16) reported that PArkin Co-Regulated Gene (PACRG) and FAP20 proteins form the IJ. PACRG and FAP20 are arranged in an alternating pattern to form the IJ linking the A- and B-tubule protofilaments A1 and B10 of the axonemal doublets. In addition, both FAP20 and PACRG are important components for motility. Both PACRG and FAP20 are conserved among organisms with cilia, suggesting a common IJ between species.

PACRG shares a bi-directional promoter with the Parkinson’s disease-related gene parkin (17, 18). Due to its axonemal functions, knockdown of PACRG genes in *Trypanosoma brucei* and *Xenopus*, lead to defects in the doublet structure and, therefore, impaired flagellar motility. In vertebrates, defects in left-right body symmetry, neural tube closure were observed from knockdowns of PACRG (19). In mice, PACRG knockout results in male sterility (20) and hydrocephalus (21). FAP20 knockout mutants in *Chlamydomonas* have motility defects and frequent splaying of the axoneme (15). Similarly, FAP20 knockdown in *Paramecium* has an altered waveform (22). A recent report identified other MIPs near the IJ, namely FAP52 and FAP45 (23). Knockouts of FAP52 or FAP45 lead to an unstable B-tubule in *Chlamydomonas*. Double knockouts of FAP52 or FAP45 together with FAP20 leads to severe damage of the B-tubule. The gene deletion of the human homolog of FAP52 has been shown to cause heterotaxy and *situs inversus totalis* in patients (24).

Cryo-EM structures of isolated doublets from *Tetrahymena* show that there are different tethering densities that connect the B-tubule to the A-tubule aside from the IJ (7, 8). However, the identity of such protein remains unknown to date. Taken together, these data suggest that there is a complex interaction at the IJ region involving multiple proteins in addition to PACRG, FAP20, FAP45 and FAP52. These interactions may play a role in regulating ciliary motility via stability.

Despite all the phenotypes known about these IJ proteins, there are no high-resolution structures to explain the molecular mechanism of the B-tubule closure and the IJ stability. In this study, we present the high-resolution cryo-EM structure of the IJ region from the *Chlamydomonas* doublet. Using a combination of sequence bioinformatics and mass spectrometry, we were able to identify two new IJ proteins, FAP276 and FAP106, and a new IJ-associated MIP, FAP126. Our results suggest that the IJ is made up of a complex of proteins involving PACRG, FAP20, FAP52, FAP276, FAP106, FAP126. We also compare the *Chlamydomonas* structure with the *Tetrahymena* structure to understand the common and species-specific features of the IJ.

## RESULTS

### Multiple tether proteins exist at the IJ

To better understand the IJ, we obtained the 48-nm repeating unit of taxol stabilized and salt treated *Chlamydomonas* doublet at 4.5 Å resolution (Fig. 1A-B and Fig. S1A-C). Due to the salt wash, some MIPs were lost when compared to the intact tomographic doublet structure (dashed parts in Fig. 1B) (5). When comparing to the cryo-EM structure of the 48-nm repeating unit of *Tetrahymena* (7, 8), the IJ region bridging PF B10 and A1 remained intact (Figure 1A-D). Based on previous studies (15, 16, 23), we were able to localize FAP52, FAP45 in both *Tetrahymena* and *Chlamydomonas* (FAP52, light green and FAP45, yellow-green in Fig. 1E, G), and PACRG and FAP20 (PACRG, light gray and FAP20, dark gray in Fig. 1F) in *Chlamydomonas*.

**Figure 1:**
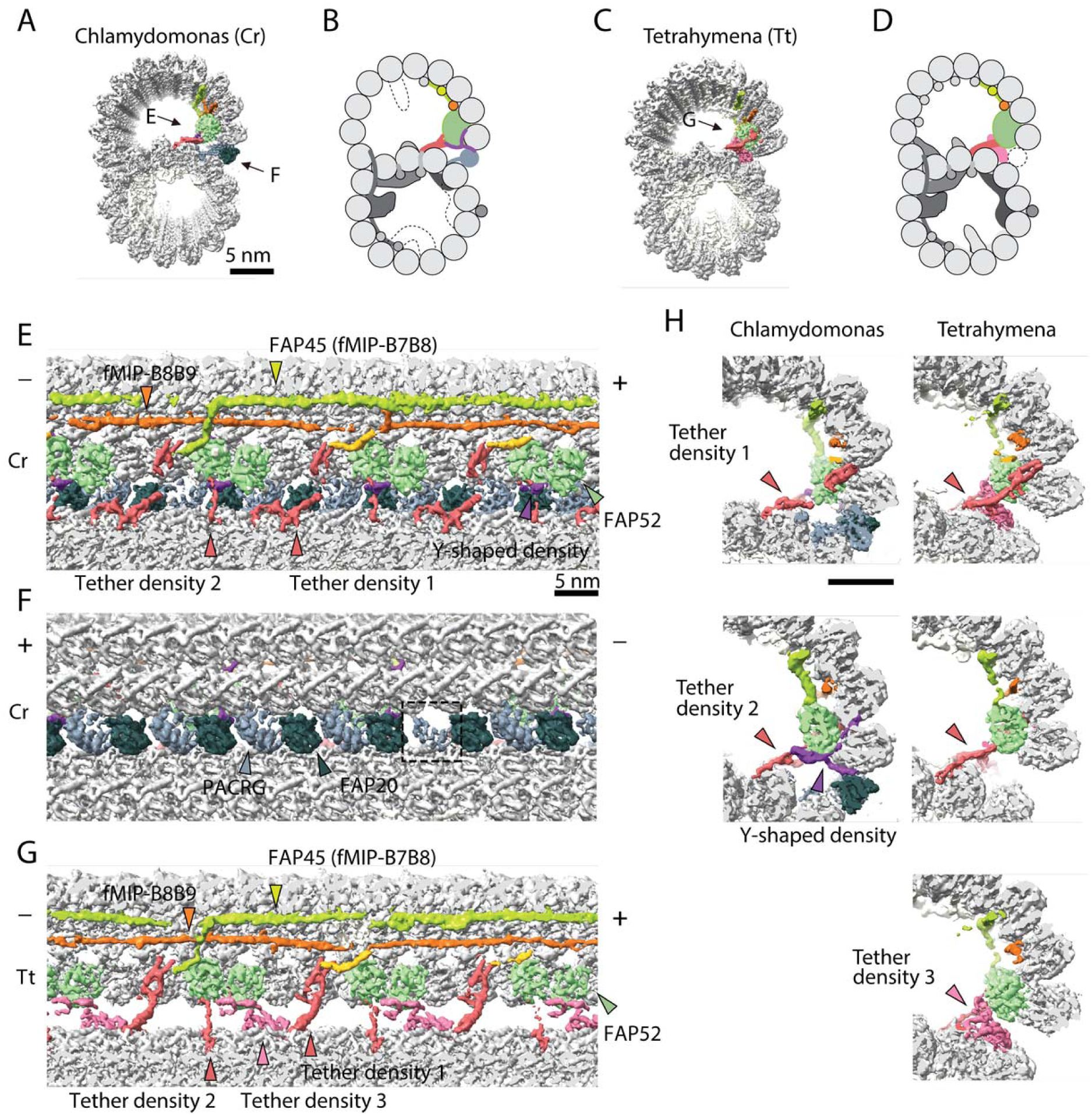
The IJ structures of *Chlamydomonas* and *Tetrahymena* doublet. (A-D) Surface renderings and schematics of the 48-nm repeat cryo-EM maps of *Chlamydomonas* (A, B) and *Tetrahymena* (C, D) doublets viewed from the tip of the cilia. Black arrow indicates longitudinal view in (E), (F) and (G). (E-F) The longitudinal section of the *Chlamydomonas* doublet at the IJ complex from the inside (E) and outside (F). (G) The longitudinal section of *Tetrahymena* doublet viewed from the inside. Color scheme: FAP20: dark gray; PACRG: gray; FAP52: light green, Y-shaped density: purple; FAP45: yellow green; fMIP-B8B9: orange; Tubulin: light gray; Rest of MIPs: white; Tether density 1 and 2: red; Tether density 3, pink. Plus and minus ends are indicated by + and - signs. (H) Cross sectional views of the different Tether densities from *Chlamydomonas* (left) and *Tetrahymena* (right). In *Chlamydomonas*, there is a Y-shaped density (purple) that cradles the FAP52 density. The Y-shaped density is absent in *Tetrahymena*. In *Tetrahymena*, we observed Tether density 3, which is absent in *Chlamydomonas*.

In this study, we termed the structure formed by the repeating units of PACRG and FAP20, the IJ protofilament (IJ PF), and refer to the IJ complex as all the proteins involved in the attachment of the B-tubule to the A-tubule. Most of the proteins in this IJ complex are attached to PFs B8 to B10 and the IJ PF.

The presence of the IJ PF stabilizes the B-tubule of the *Chlamydomonas* doublet relative to *Tetrahymena*, as evidenced by local resolution measurements (Fig. S1D). Despite having a good resolution in the A-tubule, the *Tetrahymena* doublet has a significantly lower resolution in the IJ area of the B-tubule.

Inside the B-tubule of both species, it is clear that the IJ region is held up by many tether densities along the doublet connecting the B-tubule to PF A13 (Fig. 1E-G). First, the B-tubule is held up by tether density 1 (red, Fig. 1H), referred to as MIP3b previously (7, 8). Tether density 1 connects the PF B9/B10 and A13. The second connection is named Tether density 2 (red, Fig. 1E-G), projecting from the proximal lobe of the FAP52 density (referred to as MIP3a previously (7)) and connecting to PF A13 (Fig. 1H). In *Chlamydomonas*, there is another Y-shaped density (purple) cradling the FAP52 proximal lobe density and projects into the gap between the IJ PF and PF B10 (Fig. 1H).

FAP45, which is referred to as MIP3c previously (23) is a filamentous MIP binding at the inner ridge between PF B7 and B8. In both species, FAP45 forms an L-shape density which contacts FAP52 once every 48-nm. This explains the zero-length cross-link observed in a recent study (23). In *Tetrahymena*, there exists a tether density 3 (pale violet, Fig. 1), projecting from the distal lobe of the FAP52 density and connecting to PF A13. This Tether density 3 is not present in the *Chlamydomonas* doublet, suggesting that this density is specific to Tetrahymena. All the tether densities described above repeat with 16-nm.

The IJ PF is formed by a heterodimer of PACRG and FAP20 repeating every 8-nm with the same repeating unit as tubulin dimers (Fig. 1F). This is to be expected as the purpose of the IJ PF is to bridge the tubulin dimers from PFs B10 and A1. In the 48-nm *Chlamydomonas* doublet map, however, is one PACRG unit with a less defined density compared to the others (dashed box, Fig. 1F). It has been shown that there is one PACRG density missing in every 96-nm repeat (6, 16). Since our doublet map is a 48-nm repeat unit, the less defined density of PACRG corresponds to the average from one unit of PACRG and one missing unit, i.e. half the signal. This missing unit of PACRG in the 96-nm repeat allows the basal region of the DRC to anchor onto the doublet (6) (Fig. S1F, G).

The entire IJ filament of PACRG and FAP20 seems to be missing in the *Tetrahymena* structure. Upon adjusting the threshold value of the surface rendering, we observed one dimer of PACRG and FAP20 remaining in the structure, previously named the IJ small structure (Fig. S1E) (7). This can be a result of a specific region every 96-nm of the *Tetrahymena* doublet that can hold this dimer in place preventing its detachment during sample preparation.

### PACRG, FAP20, FAP52 and FAP276 form an IJ complex

Since the majority of IJ proteins repeat with 8-nm and 16-nm, we obtained first the 16-nm repeating unit from *Chlamydomonas* and *Tetrahymena* at 3.9 Å resolution (Fig. 2A-B and Fig. S1C). Using focused refined, the IJ complex of *Chlamydomonas* was improved to 3.6 Å resolution. Without the IJ PF, the B-tubule is flexible in *Tetrahymena*, which leads to significantly lower resolution in the IJ area as shown by local resolution measurement (Fig. S1D). In contrast, the IJ region of *Chlamydomonas* has good resolution due to the stability of the B-tubule as a result of the intact IJ PF. It is worth mentioning that PFs A3-A6 in *Chlamydomonas* have lower resolution due to the lack of MIPs in this region. At 3.6 Å resolution, we were able to segment, trace and de novo model PACRG, FAP20 and FAP52 in *Chlamydomonas* (Fig. 2A, B and Fig. S2A-F). We could not model FAP45 since FAP45 repeats with 48-nm, and therefore is averaged out in the 16-nm averaged map.

**Figure 2:**
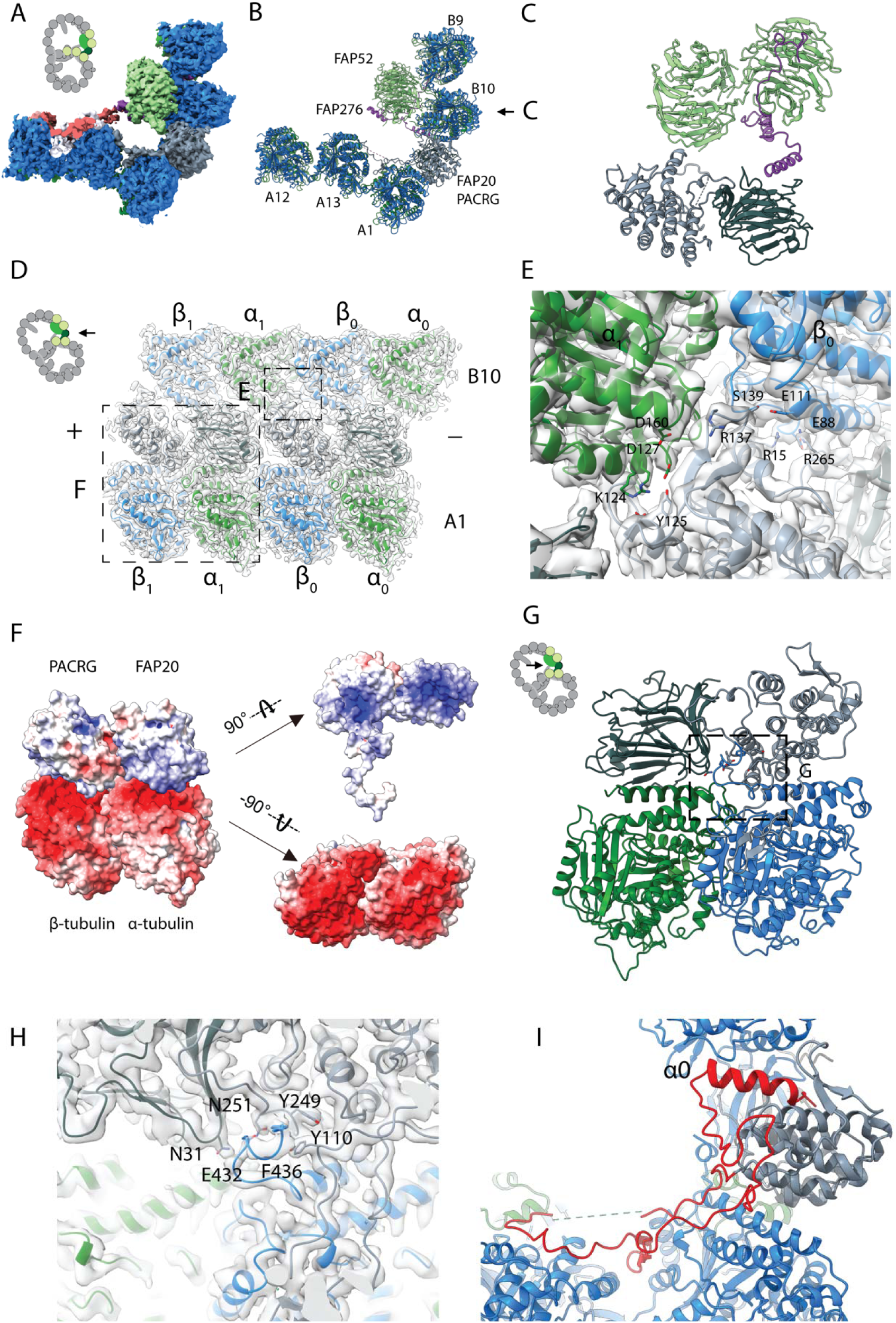
16-nm structure of *Chlamydomonas* doublet. (A, B) 16-nm repeat structure of *Chlamydomonas* doublet and model at the IJ region. (C) Atomic model of the IJ complex, consisting of PACRG, FAP20, FAP52 and FAP276. (D) Maps and model of PF A1 and B10, and IJ PF. The view is indicated in the schematic. Dashed boxes indicate the views in (E) and (F). Color scheme: α-tubulin: green; β-tubulin: blue; PACRG: gray; FAP20: dark gray; FAP276: purple. (E) The iinteraction of PACRG with the inter-dimer interface of tubulins from PF B10 is shown. (F) Electrostatic surface charge of PACRG, FAP20 and α- and β-tubulins of PF A1. Tubulin surface is negatively charged while the interacting interface of PACRG and FAP20 are positively charged. (G) The C-terminus of β-tubulin of PF A1 interacts with PACRG and FAP20. (H) Potential residues involved in the interaction of C-tail of β-tubulin and PACRG and FAP20. (I) The N-terminus of PACRG going into the wedge between PF A13 and A1. The N-terminus of *Chlamydomonas* PACRG (red color) forms a stable triple helix arrangement with the core of the protein. This is not observed in the human PACRG.

We were able to trace and, therefore, estimate the molecular weight of the Y-shaped density ∼10 kDa. Since this density is repeating with 16-nm and has a large binding interface with FAP52, we hypothesized that this protein would be missing in FAP52 knockout cells. Therefore, we did mass spectrometry of split doublets from *Chlamydomonas* FAP52 knockout cells and performed relative quantification of axonemal proteins compared to wild type (25). We observed 12 missing and 26 proteins reduced by at least 2-fold (Table 1 and Supplementary Table 1). PACRG, FAP20 and FAP45 levels are unchanged in FAP52 mutants since their binding interfaces with FAP52 are not as large as supported by our structure. The level of tektin, another suggested IJ protein does not change as well.

**Table 1:** Proteins completely missing in FAP52 knockout mutant

| **Names** | **Mol. weight**<br>**(kDa)** | **p-values**<br>**(WT vs FAP52)** | **Exclusive unique peptide counts in**<br>**WT (quantitative values after**<br>**normalization)** |
| --- | --- | --- | --- |
| ARL3 | 20 | 0.0013 | 2, 2, 1 (2, 1, 2) |
| CHLREDRAFT_171815 | 57 | 0.035 | 2, 6, 1 (2, 5, 2) |
| CHLREDRAFT_156073 | 11 | 0.024 | 1, 1, 1 (2, 1, 2) |
| FAP276 | 10 | 0.015 | 3, 3, 2 (8, 7, 14) |
| FAP52 | 66 | 0.0046 | 27, 21, 15 (59, 73, 105) |
| FAP36 | 41 | 0.0023 | 3, 3, 1 (3, 3, 2) |
| CrCDPK1 | 54 | <0.0001 | 3, 5, 2 (3, 4, 4) |
| CHLREDRAFT_176830 | 110 | 0.024 | 2, 1, 2 (2, 1, 1) |
| FAP173 | 33 | 0.012 | 3, 3, 1 (5, 3, 2) |
| FAP29 | 112 | 0.00045 | 2, 3, 2 (3, 3, 4) |
| CHLREDRAFT_181390 | 41 | 0.028 | 1, 1, 1 (1, 1, 2) |
| ANK2 | 60 | 0,0041 | 1, 2, 1 (1, 1, 2) |

Among the proteins missing in the FAP52 knockout flagella, FAP276 fits our search criteria in terms of molecular weight (Table 1). The secondary structure prediction of FAP276 and the side chains agrees unambiguously with the density signature in this region (Fig. S2G, H). This allows the unambiguous atomic modelling of FAP276 into the Y-shaped density (Fig. S2G, H). Thus, the IJ complex is made up of two copies of PACRG and FAP20, one copy of FAP52 and FAP276 and one copy of Tether density 1 and 2 per 16-nm (Fig. 2C). This represents a high stoichiometry compared to other proteins in the axoneme such as CCDC39 and CCDC40, which have only one copy per 96-nm (26).

The PACRG structure is composed mainly of α-helices with a long unstructured N-terminal region. PACRG contains an alpha solenoid architecture, similar to the microtubule binding TOG domain, which is present in many microtubule polymerases (27, 28). On the other hand, FAP20 has a beta jelly roll architecture, which consists of mainly β-sheets with a small α-helix. The C-terminus of FAP20 is located at the outside of the doublet, in agreement with a tomographic study of FAP20 with a Biotin Carboxyl Carrier Protein tag at the C-terminus. (15).

PACRG and FAP20 have two microtubule-binding sites, one on the surface of the A-tubule similar to well-studied microtubule-associated protein binding sites such as TOG (28), and one on the lateral side of the B-tubule (Fig. 2D). The lateral binding site is unique and has never been observed in previously known microtubule-associated proteins. PACRG binds to the inter-dimer interface of PF B10 in the region of MEIG1 binding loop (29) and β-tubulin from PF A1. The interaction with PF B10 involves residues Y125, R137, S139 and R265 with E88 and E111 from β-tubulin and D160 and D127 from α-tubulin (Fig. 2E). FAP20 is sandwiched by the tubulin dimer from PF B10 and the α-tubulin from PF A1 (Fig. 2D).

The interactions of PACRG and FAP20 with tubulin from PF A1 appear to be electrostatic. The outside surfaces of α- and β-tubulins are highly negatively charged while the corresponding interacting surfaces of PACRG and FAP20 are positively charged (Fig. 2F). Despite the fact that PACRG contains alpha solenoid architecture like TOG domains, the binding orientation of PACRG to the surface of tubulin is completely different from TOG domain binding (28).

In addition to the interactions highlighted above, we also observed the interaction of the β-tubulin C-terminus from PF A1 with PACRG (Fig. 2G, H). The C-termini of α- and β-tubulins are a hot spot for post-translational modifications such as polyglutamylation and polyglycylation (30). However, due to its flexibility, densities for the α- and β-tubulin C-termini are usually not visible in microtubule cryo-EM reconstructions. This is also the case for the outside of the A- and B-tubules in our ex vivo structure. However, in the lumen of the B-tubule, the β-tubulin C-terminus from PF A1 appears to be stabilized by two key interactions with PACRG: the hydrogen bond between D432 together with the hydrophobic burial and the stable T-shaped stacking of F436 with N251 and Y249 of PACRG, respectively (Fig. 2H). Both interactions stabilize the β-tubulin C-terminus forming a helical turn in segment E432-E437, which otherwise wouldn’t be present due to its flexibility.

This structuring of the β-tubulin C-terminus in PF A1 appears to be the result of the steric proximity with the N-terminus of PACRG. Thus, both interactions are important in maintaining the stability of the IJ by preventing steric clashing between the two. It could also be an indication of further post-translational modifications that occur in this region, which could have a potential role in IJ formation and stability.

In our structure, we also observe that the distance between FAP20 and the proximal PACRG is closer compared to the distal PACRG, thus PACRG and FAP20 likely form a heterodimer instead of a continuous protofilament (Fig. 3A). The PACRG and FAP20 binding interface appears to involve multiple hydrogen bonds with complementary surface charges, suggesting a specific and strong interaction (Fig. 3B, C). This FAP20 binding loop of PACRG is well-conserved among species (Fig. 3D). This loop is not presented in the PACRG-like protein, a homolog of PACRG and exists in the basal body (29). FAP20, on the other hand, has a high degree of sequence conservation (Fig. S3).

**Figure 3:**
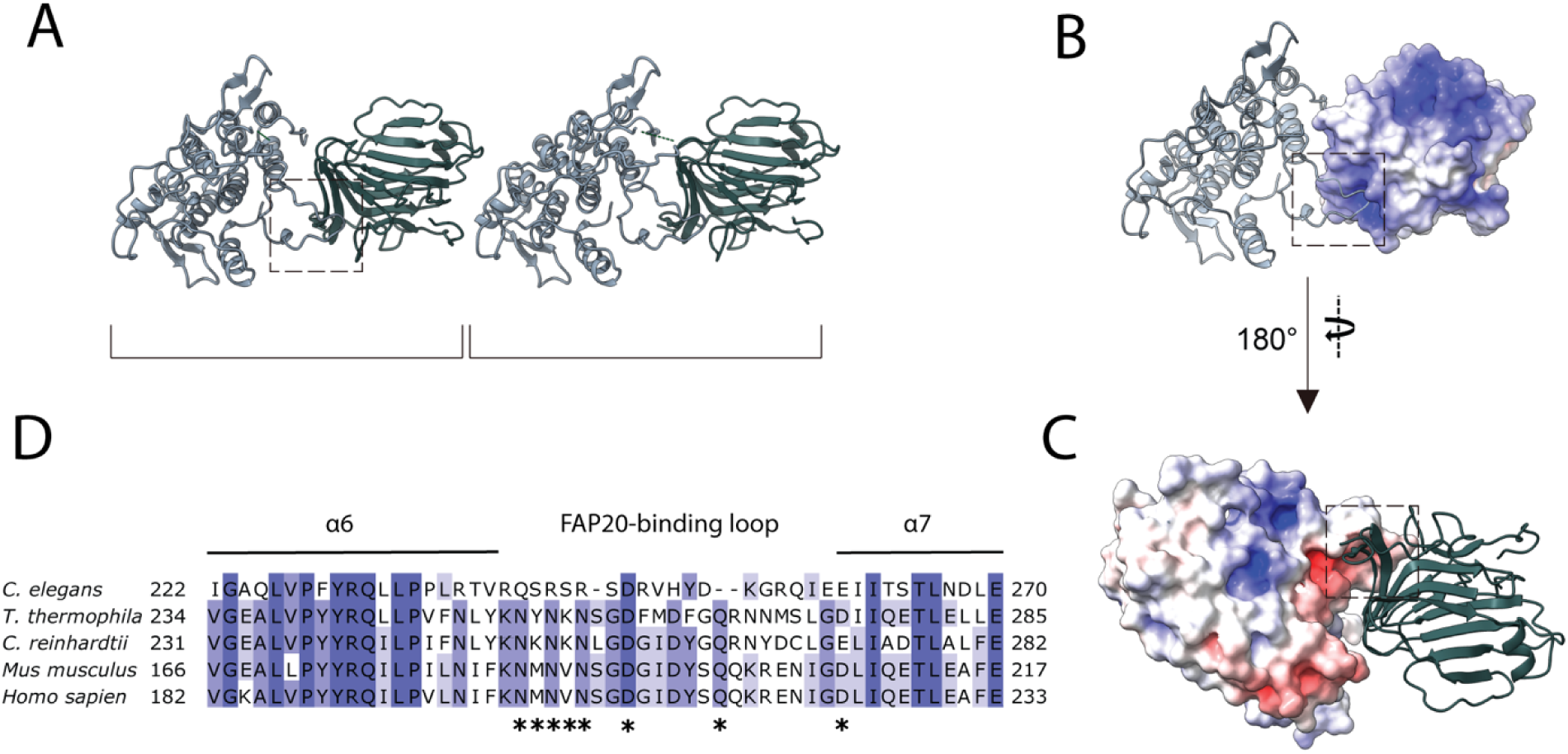
PACRG and FAP20 form a homo dimer. (A) Consecutive molecules of PACRG and FAP20 in the IJ protofilament. PACRG and FAP20 form a heterodimer as indicated by brackets. (B, C) Electrostatic interactions between PACRG and FAP20 illustrated by their surface charge. The dashed boxes in (A, B, C) highlight the interacting loops between PACRG and FAP20. (D) Multiple sequence alignment of PACRG in the regions of FAP20-binding loop. Asterisks indicate residues that are involved in FAP20 binding.

The cryo-EM structure of the *Chlamydomonas* PACRG is highly similar to the crystal structure of the human PACRG binding to MEIG1 (PDB: 6NDU) (29), suggesting a conserved role of PACRG. *Chlamydomonas* PACRG has a long N-terminus that binds on top of PF A13 and into the wedge between PF A1 and A13 (Fig. 2I, Fig. S2A). This N-terminal region is not conserved in humans or *Tetrahymena* (29). This could indicate organism-specific adaptations to achieve finely tuned degrees of ciliary stability.

### FAP52 forms an interaction hub and stabilizes α-tubulin’s acetylated K40 loop

Next, we investigated the structure of FAP52 (Fig. S2C). FAP52 consists of eight WD40 domains forming two seven-bladed beta-propellers. The two beta-propellers form a V-shape that docks onto PF B10 and B9. The proximal beta-propeller docks onto the inside of the α- and β-tubulin intra-dimer interface, while the distal beta-propeller is aligned with the next inter-dimer interface towards the plus end (Fig. 4A).

**Figure 4:**
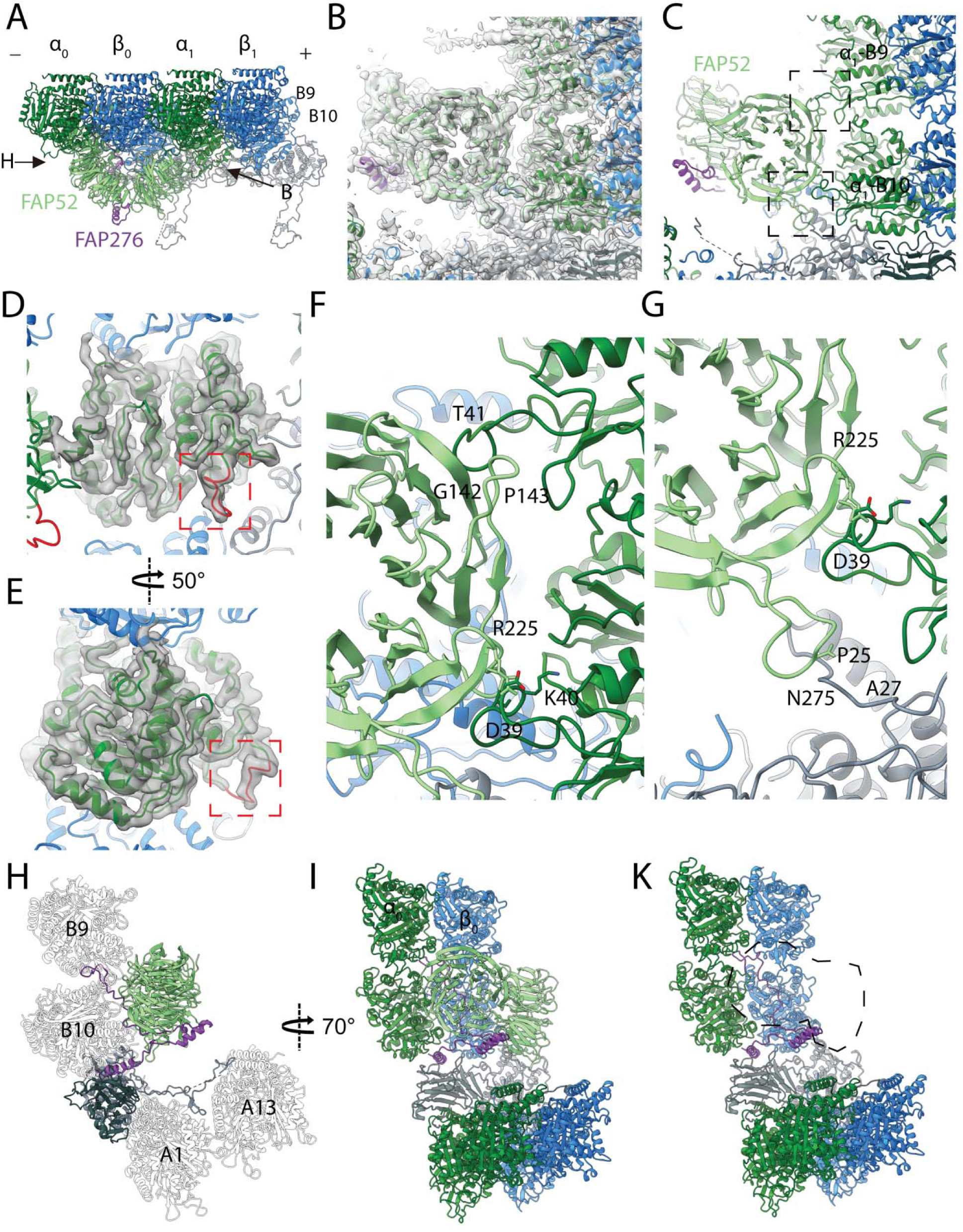
Structure of FAP52 and its interaction with tubulins and PACRG. (A) Structure of FAP52 in a top view from the outside of the B-tubule looking down on the A-tubule. Black arrows indicate the direction of view in (B) and (H). (B, C) Interactions of FAP52 with α-tubulins from PF B8 and B9 and PACRG with (B) and without map overlay (C). (D-E) The structure of the α-K40 loop from PF B10. Red dashed boxes indicate the α-K40 loop. (F) Interaction of α-K40 loop of PF B9 and B10 with FAP52. In PF B10, residue D39 of α-tubulin appears to form a salt bridge with R225 from FAP52. In PF B9, T41 of α-tubulin appears to interact with segment G142-P143 from FAP52. (G) N275 from loop V268-L279 of FAP52 appears to interact with the N-terminus of PACRG in segment P25-A27. Loop V268-L279 in *Chlamydomonas* FAP52 is missing in the *Tetrahymena* and the human structures. (H-K) Interactions of FAP276 with FAP52, FAP20 and tubulins. In (K) FAP52 model is digitally removed to show the interactions underneath.

The distal beta-propeller of FAP52 has a 3-point contact with the inner surface of the B-tubule (Fig. 4B, C). Two of the FAP52 contacts involve the K40 loop of α-tubulin from PF B9 and B10. The α-K40 acetylation was first discovered in *Chlamydomonas* flagella, which is almost fully acetylated (31). This α-K40 loop has not been fully visualized in reconstituted studies of acetylated tubulins. In our structure, the α-K40 loop is fully structured in this position (Fig. 4D, E and Fig. S4D-G). For the first contact point, residue R225 of FAP52 seems to interact with D39 of α-tubulin from PF B10 (Fig. 4F, G). At the second tubulin contact point, FAP52 segment G142-P143 appears to interact with T41 of α-tubulin from PF B9. In the lower region of FAP52, residue N275 from the distal beta-propeller’s V268-L279 loop interacts with segment P25-A27 of the N-terminus of PACRG (Fig. 4G). The density of the aforementioned loop is not present in the FAP52 structure in Tetrahymena doublet (Fig. S4A, B). This long loop (V268 to L279) of *Chlamydomonas* FAP52 is, in fact, deleted in other species (Fig. S4C). The interaction of this loop with *Chlamydomonas* PACRG suggests that it is a *Chlamydomonas* specific feature that stabilizes PACRG and, hence the IJ PF.

We then investigated the α-K40 loops from *Chlamydomonas* and *Tetrahymena* doublets (Fig. S4D-G). When there is no interacting protein, this loop is flexible consistent with previous literature (32). Despite having low resolution in the B-tubule in *Tetrahymena*, we still observed the α-K40 loop of PF B9 and B10 interacts with FAP52 (Fig. S4B). We were also able to visualize the loop in several places in both *Chlamydomonas* and *Tetrahymena* where there is an interacting protein (Fig. S4F, G). The conformation of the loop appeared to be different depending on its interacting protein. This suggests that the α-K40 loop could have a role in MIP recognition and binding.

Furthermore, because of the V-shape of FAP52, its interacting interface with tubulin is small. The existence of a cradling protein such as FAP276 then is logical from a functional standpoint since it appears to support and mediate the interaction between FAP52 and tubulin (Fig. 4H). Segment L52-H57 from FAP276 forms beta sheet-like stacking interactions with segment L375-V380 from FAP52 (Fig. 4H, I). FAP276 itself forms numerous interactions with tubulin with both of its N- and C-termini, thus it provides strong anchorage for FAP52 to the tubulin lattice (Fig. 4H-K). Given the numerous interactions of FAP52 with all the proteins mentioned, FAP52 is likely to function as an interaction hub, which could play an important role during IJ assembly.

### FAP106 is the Tether loop, consisting of Tether density 1 and 2

We were able to trace the Tether density in the 16-nm averaged map. Tether density 1 is connected to Tether density 2 (Fig. 5A-D), forming a Tether loop, through which the A- and B-tubules are connected. The loop connecting the Tether density 1 binds on top of PF A12 and then into the outside wedge between A12 and A13 before connecting with Tether density 2. Therefore, the entire Tether loop is a single polypeptide, conserved between *Tetrahymena* and *Chlamydomonas* (Fig. 5A-B). Part of this Tether loop resembles Tau binding to the microtubule (33). There is a small helical region in this loop that binds to α-tubulin of PF A12 (Fig. S5A, B).

**Figure 5:**
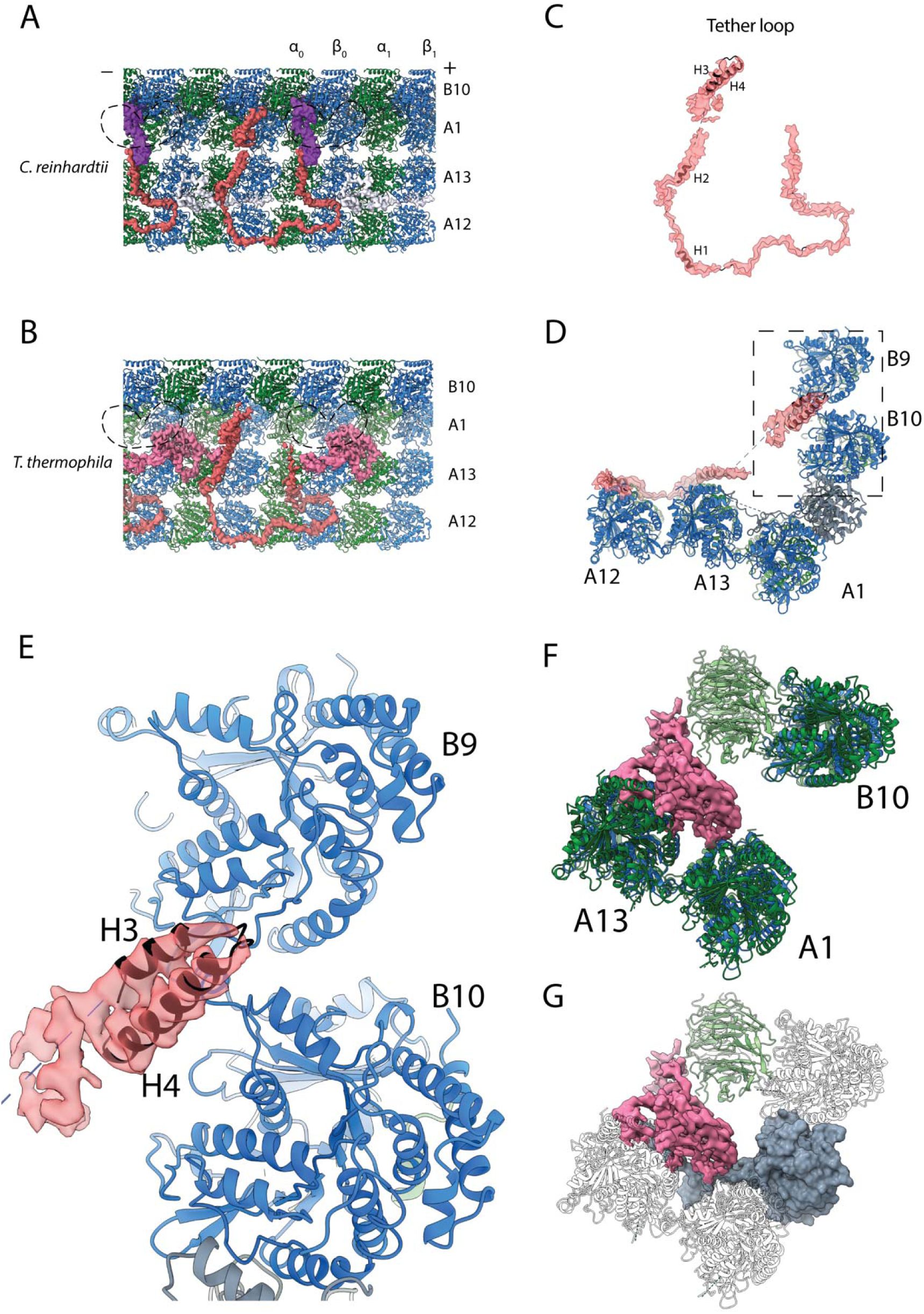
Structure of the Tether densities. (A-B) At higher resolution, Tether densities 1 and 2 appear to be a single polypeptide chain in both *Chlamydomonas* (A) and *Tetrahymena* (B). The dashed regions indicate the location of FAP52, which has been digitally removed to show the Tether densities underneath. (C) Model of FAP106 fitted inside the segmented Tether loop from *Chlamydomonas*. (D) Model of FAP106 tethering the B-tubule and A-tubule. Dashed box indicates view in (E). (E) Helix H3 and H4 of FAP106 insert into the gap formed by four tubulin dimers of PF B9 and B10. (F) Structure of Tether density 3 from *Tetrahymena*, which binds on top of the wedge between PF A13 and A1. (G) Overlay of the PACRG from *Chlamydomonas* onto the structure of *Tetrahymena* shows a hypothetical steric clash of a long *Tetrahymena* PACRG N-terminus with Tether density 3.

To identify the protein that makes up the Tether loop, the protein needs to satisfy the following criteria: (i) has a high stoichiometry (1 per 16-nm of the doublet); (ii) has a minimum molecular weight of ∼25kD (based on a poly-Alanine trace) and (iii) conserved in both *Chlamydomonas* and *Tetrahymena*.

We calculated the stoichiometry of proteins in the doublet after salt extraction by normalizing the averaged quantitative spectral count of each protein by their molecular weight. The triplicate mass spectrometry data comes from Dai et al. (25). The top 35 proteins by copy numbers are shown in Table 2. In our calculation, some radial spoke and central pair proteins displayed high stoichiometry such as RSP9 and PF16. Remarkably, all the IJ proteins are in the top 35 (PACRG, FAP52, FAP20, FAP45 ranked 4, 9, 10 and 35 respectively) as supported by our structure. This validates the quality of the stoichiometry calculation. Although FAP276 should have the same stoichiometry as FAP52, it does not appear in high stochiometric numbers. This can be explained that by the small size of FAP276, which is not well detected in mass spectrometry.

**Table 2:** Normalized spectral count of proteins detected by mass spectrometry.

| Name | Molecular Weight in kDa (MW) | Average Quantitative Value (AQV) | Rough Stoichiometric Peptide Abundance (RSPA)* | *T. thermophila* Homologs** | Human Homologs*** | Localization in *C. reinhardtii* |
| --- | --- | --- | --- | --- | --- | --- |
| TUA1 | 50 | 1077.97 | 215.59 | TBA_TETTH | TUBA1C | Doublet |
| TUB1 | 50 | 625.46 | 125.09 | TBB_TETTH | TUBB4B | Doublet |
| RIB72 | 72 | 116.72 | 16.21 | TTHERM_00143690 | EFHC1 | MIP |
| PACRG | 25 | 38.39 | 15.35 | TTHERM_00446290 | PACRG | IJ |
| PF16 | 50 | 74.09 | 14.82 | TTHERM_000157929 | SPAG6 | Central Pair |
| RSP9 | 30 | 41.79 | 13.93 | TTHERM_00430020 | RSPH9 | Radial Spoke |
| FAP86 | 30 | 36.11 | 12.04 | - | - | Doublet |
| FAP1 | 22 | 26.46 | 12.03 | - | - | Doublet |
| FAP52 | 66 | 79.12 | 11.99 | TTHERM_01094880 | CFAP52 | MIP |
| FAP20 | 22 | 26.08 | 11.86 | TTHERM_00418580 | CFAP20 | IJ |
| FAP126 | 15 | 16.89 | 11.26 | - | CFAP126 | MIP |
| RSP1 | 88 | 98.73 | 11.22 | TTHERM_00047490 | RSPH1 | Radial Spoke |
| FAP115 | 27 | 29.98 | 11.10 | TTHERM_00193760 | - | Doublet |
| FAP106 | 27 | 29.81 | 11.04 | TTHERM_00137550 | ENKUR | IJ? |
| Tektin | 53 | 57.61 | 10.87 | - | TEKT5 | IJ? |
| RSP3 | 57 | 60.55 | 10.62 | TTHERM_00566810 | RSPH3 | Radial Spoke |
| FAP252 | 39 | 39.97 | 10.25 | TTHERM_00899430 | CETN3 | Axonemal |
| RSP2 | 77 | 77.95 | 10.12 | - | CALM2 | Radial Spoke |
| FAP161 | 43 | 43.50 | 10.12 | TTHERM_00155380 | CFAP161 | Axonemal |
| IDA4 | 29 | 27.29 | 9.41 | TTHERM_00841210 | DNALI1 | Dynein |
| FAP107 | 26 | 23.66 | 9.10 | - | FLG2 | Axonemal |
| DHC2 | 457 | 414.12 | 9.06 | TTHERM_01027670 | DNAH1 | Dynein |
| FAP12 | 54 | 48.85 | 9.05 | - | DAGLB | Cytoplasmic |
| RSP7 | 34 | 30.62 | 9.01 | TTHERM_00194419 | CALML5 | Radial Spoke |
| RSP5 | 56 | 49.32 | 8.81 | - | - | Radial Spoke |
| FAP230 | 45 | 39.59 | 8.80 | - | - | Axonemal |
| FAP77 | 29 | 23.88 | 8.24 | TTHERM_00974270 | CFAP77 | Axonemal |
| FAP55 | 111 | 90.47 | 8.15 | - | MYH14 | Axonemal |
| FAP90 | 28 | 22.35 | 7.98 | - | WBP11 | Axonemal |
| RSP10 | 24 | 19.10 | 7.96 | TTHERM_00378600 | RSPH1 | Radial Spoke |
| FAP71 | 32 | 24.86 | 7.77 | TTHERM_00077710 | EWSR1 | Axonemal |
| EEF1 | 51 | 39.04 | 7.66 | TTHERM_00655820 | Multiple | Axonemal |
| FAP182 | 49 | 36.69 | 7.49 | TTHERM_01049330 | C9orf116 | Axonemal |
|  |  |  |  | TTHERM_00624660 | RIBC2 | MIP |
| Rib43a | 43 | 32.06 | 7.46 | TTHERM_00641119 |  |  |
| FAP45 | 59 | 43.20 | 7.32 | TTHERM_001164064 | CFAP45 | MIP |

\*RSPA was calculated by (AQV)/(MW)\*10
\*\* *T. thermophila* homologs were BLASTed using the Uniprot database
\*\*\* Human homologs were taken from the ChlamyFP project

Among the proteins that have high stoichiometry, the following proteins satisfy the three criteria above: FAP115, FAP106, FAP252, FAP161, FAP77 and FAP71. However, the homologs of FAP115 and FAP161 in *Tetrahymena* are too big. Our analysis of the secondary structure prediction places FAP106 at the top of the list of candidates for the Tether loop (Fig. S5D). Furthermore, the sequence agrees with the density signature unambiguously, which leaves no doubt that the Tether loop is FAP106. This allowed us to model segments P20-A148 and W189-I226 where the density had sufficient signal (Fig. 5C, D). Segments M1-R19 and R227-D240 have low SNR and are likely to be highly flexible. Helix H3 and H4 of FAP106 insert into the interdimer interfaces between PF B9 and B10 forming the anchor point to the B-tubule (Fig. 5E) while helix H1 and H2 bind to β-tubulin of PF A13 and α-tubulin of PF A12 (Fig. 5D). FAP106 is a homolog of ENKURIN (ENKUR), a conserved protein in sperms of many species (34, 35). Enkur knockout mice have abnormal sperm motility with asymmetric flagellar waveform and therefore low fertility rate (35). In addition, mutations in ENKUR is linked to situs inversus in human and mouse (36, 37). However, the IQ motif of Enkurin that binds Calmodulin is not conserved in Chlamydomonas (Fig. S4C).

In *Tetrahymena*, the Tether density 3 connects the distal lobe of FAP52 and binds across the wedge between PF A13 and A1. (Fig. 5F). Upon superimposing the *Chlamydomonas* PACRG structure onto the *Tetrahymena* IJ area, the N-terminus of PACRG will have a steric clash with Tether density 3 (Fig. 5G). This explains the shorter N-terminus of *Tetrahymena* PACRG relative to the *Chlamydomonas* PACRG. Tether density 3 might interact with and perform the same function as the N-terminus of PACRG in *Chlamydomonas*, which induces high curvature of PF A13 and A1.

### FAP126, a FLTOP homolog, interacts with the tether loop, FAP106

We also were able to trace a density that lies on top of PF A13 and goes into the wedge between PF A12 and A13 (Fig. 6A, turquoise). This density was described previously as part of MIP5 (7), and is mostly disordered. It is not present in the *Tetrahymena* map and was traced as a single polypeptide.

**Figure 6:**
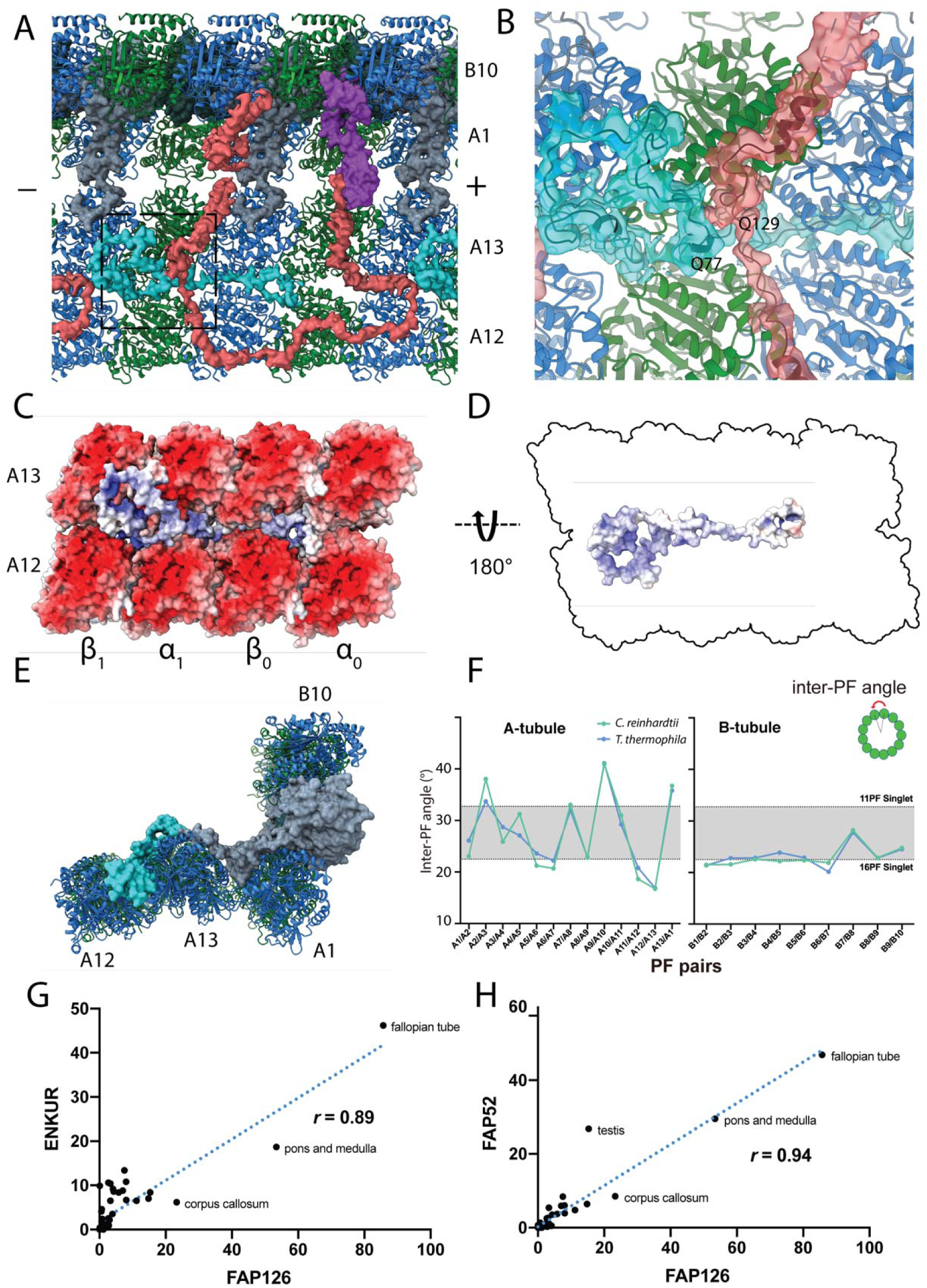
Structure of FAP126 and its interaction. (A) View from the top of the PF A12 and A13 showing the density of FAP126 (dark turquoise). Dashed box indicates view in (B). (B) Close up view of FAP126’s interaction with the Tether loop, FAP106 at residues N74-T76 and Q129. (C) Complimentary electrostatic surface charges of tubulins and FAP126. (D) Electrostatic charge of FAP126 on the tubulin interacting surface. (E) The N-terminus of PACRG and the hook density go into the wedges between PF A12 and A13, and PF A13 and A1, respectively. This likely contributes to the curvature of this region. (F) Inter-PF angles of the A- and B-tubules from *Chlamydomonas* and *Tetrahymena* showing very similar angle distributions. (G) and (H) Correlation graphs of consensus normalized expression levels for two selected pairs of genes (ENKURIN(FAP106)/FAP126 and FAP52/FAP126). Tissues showing high levels of expression of one or both genes are labeled. Correlation coefficients (*r*) are indicated.

The density in this region had multiple clear side chains that could be identified, and so we applied two search strategies to identify this protein. We used a local search against the entire proteome of *Chlamydomonas* using a regular expression that matches a pattern of WPxxxxxW which was observed in the density. This resulted in one unique hit, which is FAP126. We then applied the same strategy to search for this protein as FAP106. The criteria were: (i) a high stoichiometry number; (ii) a size of ∼15 kDa and (iii) no homolog in *Tetrahymena*. In this case, the only protein that satisfied these criteria among high stoichiometry proteins (Table 2) was also FAP126. Furthermore, the remainder of the FAP126 sequence agrees unambiguously with the density signature in this region (Fig. 6B and Fig. S5B).

FAP126 is a homology of human FLTOP, a protein known for basal body docking and positioning in mono- and multi-ciliated cells (38). Multiple alignment sequence alignment of FAP126 shows that the *Chlamydomonas* FAP126 lacks the proline-rich regions of other species (Fig. S5A).

FAP126 appears to interact with FAP106 (Fig. 6B). Segment F75-Q77 of FAP126 is in proximity to segment T128-K130 of FAP106. Q77 and Q129 of FAP126 and FAP106, respectively, are within favorable distance and orientation to form a hydrogen bond with one another (Fig. 6B). Therefore, FAP126 might play a role in recruiting FAP106 or vice versa. Almost half of FAP126 density runs along the wedge between PF A12 and A13, close to the tubulin lateral interface with complementary surface charge (Fig. 6C, D). FAP126 might act as a low curvature inducer from the outside similar to Rib43a from the inside since the curvature of A12 and A13 is significantly lower compared to 13-PF singlet (Fig. 6E, F) (8).

To support whether FAP126 interacts with FAP106, we analysed the normalized RNA expression of FAP126 with FAP106 (ENKUR) and FAP52 from different human tissues (Fig. 6G, H and Fig. S5C). FAP126 showed high correlation with both FAP106 (ENKUR) and FAP52(r=0.89, p-value =<0.0001, r=0.94, p-value=<0.0001, respectively). This indicates that FAP126 might be functionally related to other members of the IJ such as FAP106 and FAP52, which further supports the identity of these proteins.

## DISCUSSION

In this study, we describe the complete molecular details of the IJ complex using a combination of mass spectrometry and cryo-EM. The IJ complex in *Chlamydomonas* is made up of PACRG, FAP20, FAP52, FAP276, FAP106 (Tether loop) and associated proteins such as FAP45 and FAP126 (Fig. 7A). We identified two new members of the IJ, FAP106 and FAP276. FAP276, a *Chlamydomonas* specific protein, anchors and mediates FAP52 onto tubulins from PF B9 and B10. FAP106 tethers the B-tubule to the A-tubule, through its interactions with the PF A12 and A13, FAP52 and FAP276, while the IJ PF, composed of PACRG and FAP20, closes the IJ gap. For the doublet to withstand the mechanical strain during ciliary beating, it needs all its unique structural features and interactions for proper stability. Tektin, a coiled-coil protein, was also proposed to be another component of the IJ complex in *Chlamydomonas* by biochemical experiments (15). However, no filamentous density corresponding to tektin was found at the IJ PF in our *Chlamydomonas* map.

**Figure 7:**
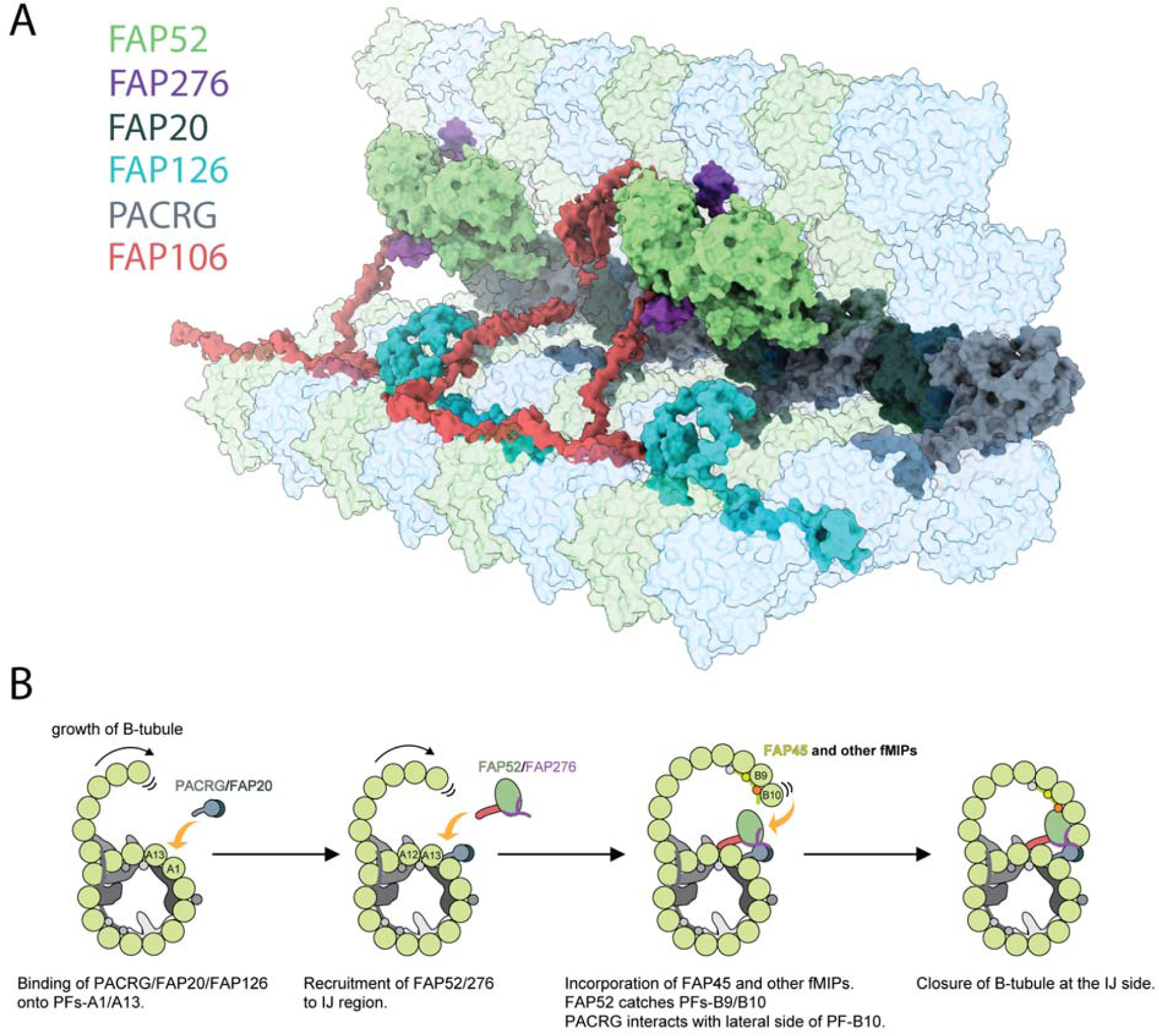
Proposed mechanism of IJ formation and B-tubule closure. (A) Model of the IJ complex including PACRG, FAP20, FAP52, FAP126, FAP276 and the Tether loop. FAP45 is not depicted here. Tubulin is depicted as transparent. (B) The B-tubule starts growing laterally from the outer junction side as shown in (14). PACRG and FAP20 form a hetero-dimer, which binds onto the outside surface of PF A1. After which, multiple alternative hypotheses are possible. One hypothesis is that FAP52, FAP276 and the Tether density proteins would bind onto PF A12 and A13. FAP45 and other fMIP proteins would then be incorporated inside the B-tubule, which fixes the proper curvature so that PF B9 and B10 can interact with other IJ proteins. FAP52 binds both PF B9 and B10 through their K40 loops and finally, PACRG and FAP20 interact with the lateral side of PF B10 allowing for B-tubule closure.

Reconstituted doublet microtubules (14) indicate that the B-tubule cannot be closed and is extremely flexible without the IJ PF. Therefore, the IJ PF is necessary to dock the B-tubule onto the A-tubule. In our *Tetrahymena* doublet, in which the IJ PF was washed away, even with the presence of FAP52 and FAP106, the doublet is still flexible which can be seen by the lower resolution of the B-tubule compared to the A-tubule (Fig. S1D). In addition, the B-tubule can be subjected to depolymerization when the IJ PF is not fully formed (23). Therefore, the IJ PF serves as an anchor, which protects the B-tubule from depolymerization by shielding the lateral side of PF B10 (Fig. 7B). Because of the complexity of interactions and the diverse protein composition of the IJ complex, it is reasonable to assume that the IJ is assembled after the outer junction nucleates and expands towards the IJ. The IJ complex might be assembled or co-assembled at the same time as PF B10 for the closure of the B-tubule (Fig. 7B).

During doublet assembly, unorderly binding of PACRG and FAP20 to any of the PFs in the B-tubule lateral interfaces would lead to an incomplete B-tubule (29). To facilitate a successful IJ assembly, chaperones might be needed for the transport of PACRG and FAP20. PACRG forms a complex with MEIG1 (29). Even though MEIG1 is not present in lower eukaryotes and that the MEIG1 binding loop is not conserved between *Chlamydomonas* and humans, a chaperone similar to MEIG1 can function to target PACRG to the lateral interface of the PF B10. Our atomic models support that PACRG and FAP20 might form a heterodimer before their transport and assembly into the cilia (Fig. 3A). Surprisingly, FAP20 shows a similar fold and mode of binding to a class of proteins called carbohydrate-binding modules. Carbohydrate binding modules form a complex with carbohydrate-active enzymes and are known to have a substrate targeting and enzyme-concentrating function (39). This supports the role of FAP20 as an assembly chaperone in a FAP20-PACRG complex. Furthermore, both studies from Yanagisawa et al. (15) and Dymek et al. (16) show reduced endogenous PACRG in *Chlamydomonas* FAP20 knockout mutant. In the latter study, it was shown that the assembly of exogenous PACRG was less efficient in the FAP20 knockout compared to conditions where FAP20 was intact. This implies that PACRG assembly might indeed depend on FAP20 (15). However, since the expression patterns of PACRG and FAP20 have surprisingly low correlation (Fig. S5E) compared to the rest of the IJ proteins, it suggests that FAP20 might have an additional function outside the IJ of the cilia.

Furthermore, our atomic models could explain the severe motility phenotypes observed in PACRG and FAP20 mutants compared to FAP52 mutant. Mutants in either PACRG or FAP20 might affect the stability of the DRC, which can severely affect the regulation of ciliary beating. This is supported by the fact that FAP20 mutant is prone to splaying of the cilia (15). Our results could also explain how the double knockout of FAP20 along with FAP45 or FAP52 can affect B-tubule stability at the IJ (23). In such conditions, both the IJ PF and the FAP52 or FAP52-mediated anchorage between the A- and B-tubules will be completely lost.

By comparing *Chlamydomonas* and *Tetrahymena*, we show that the conserved IJ components are PACRG, FAP20, FAP45, FAP52 and FAP106. There are also species-specific proteins such as FAP276 and FAP126 in *Chlamydomonas* and the Tether density 3 in *Tetrahymena*. The Tether density 3 clashes with a superimposed *Chlamydomonas* FAP276 structure, suggesting that it takes over its role in mediating the interactions between FAP52 and tubulin in *Tetrahymena*. This suggests that there is a common framework for the IJ complex in all species. Species-specific proteins may then fine-tune this framework according to the organism’s survival needs.

In this study, we revealed that FAP106/ENKUR, an important protein for sperm motility, is a MIP and inner junction protein. Knockout of ENKUR leads to the asymmetric waveform of sperm flagella while mutations in ENKUR disturb the left-right symmetry axes in vertebrates. It is shown that knockout ENKUR shows a loss of Ca^++^ responsiveness while wild type sperm shows highly curved flagella (35). The IQ domain responsible for Ca^++^ binding of ENKUR is not conserved in the Chlamydomonas sequence, although an alternative means of Ca^++^ binding or inducing a Ca^++^ mediated response is still possible.

In this study, we also identified FAP126, an inner junction-associated MIP. The homolog of FAP126 in human and mouse, the FLTOP protein, exists in the cilia and basal bodies and functions to position basal bodies (38). In Flattop knockout mice, cilia formation in the lung is significantly affected. In the inner ear, Flattop interacts with a protein called Dlg3 in the process of basal body positioning to the actin skeleton in the inner ear. Therefore, it is possible that FAP126 might perform both functions (i) as a MIP that stabilizes the basal bodies and cilia and (ii) as a basal body positioning and planar cell polarity complex. The high correlation of FAP126 and FAP106 co-expression, and also with FAP52 in different human tissue suggests they might function similarly or co-operatively in the cilia assembly. *Tetrahymena*, which lacks FAP126, probably uses an alternative mechanism for its basal body positioning.

Our study demonstrates that all the protein components associated with the IJ complex such as PACRG, FAP20, FAP52, FAP126 and FAP106 in the IJ complex is of high importance for the assembly and proper motility of the cilia. Multiple studies have indeed showed the implication of such proteins in human disease (20, 24, 37). Remarkably, these proteins are MIPs existing inside the doublet except for PACRG and FAP20. This revelation supports the notion that MIPs can directly influence the tubulin lattice and hence the motility of the cilia (8). In the regions, where MIPs are not present due to preparation, the local resolution is significantly worse than the global resolution (Fig. S1D). In addition, the structures of the IJ components such as PACRG and FAP126 also highlight the unique roles of the MIPs in curvature inducing or sensing as shown previously with Rib43a (8). FAP126 binds tightly to the wedge PF A12 and A13 while the N-terminus of *Chlamydomonas* PACRG penetrates the wedge between PF A13 and A1 and forces the PF pairs into a high-curvature conformation (Fig. 6E). From our curvature analysis of the doublet (Fig. 6F), this region of PF A12-A1 contains extreme high and low curvatures compared to the 13-PF singlet. Alternatively, the curvature might be enforced by MIPs inside the A-tubule. This inter-PF curvature could help to facilitate the specific binding and anchoring of FAP126 and PACRG to the right position. It has been shown that doublecortin can sense the curvature of the 13-PF microtubule (40). In *Tetrahymena*, Tether density 3, which is not present in *Chlamydomonas* seems to be a high curvature inducer/sensor and an IJ complex stabilizer.

Post-translational modifications in tubulin are known to be important for the activity of the cilia. There have been many studies about the effect of acetylation on the properties of microtubule such as stability (41, 42). In 3T3 cells, the K40 acetyltransferase, αTAT1 promotes rapid ciliogenesis (43). The absence of acetylating enzymes has indeed been shown to affect sperm motility in mice (44) while SIRT2 deacetylation decreases axonemal motility in vitro (45). A recent cryo-EM study of reconstituted acetylated microtubules showed, using molecular dynamics, that the acetylated α-K40 loop has less conformational flexibility, but a full α-K40 loop in the cryo-EM map has not been visualized due to its flexibility. In this work, we show that the acetylated K40 loop binds to FAP52 and forms a fully structured loop. This loop remains flexible and unstructured when there is no interacting protein. This suggests that the α-K40 loop has a role in protein recruitment and interactions, especially, MIPs. We hypothesize that the acetylation disrupts the formation of an intra-molecular salt bridge between K40 and D39, which affects the loop’s sampling conformations and allows D39 to take part in atomic interactions with other proteins. This, in turn, improves the stability of the doublet and therefore, correlates with axonemal motility. In neurons, microtubules are also highly acetylated and are known to be stable. Our hypothesis suggests that in neuron microtubules, there might exist MIPs with a similar stabilizing effect as in the doublet. Previous studies on olfactory neurons demonstrate that there are densities of proteins inside the microtubule, suggesting the existence of MIPs inside cytoplasmic microtubules (46).

Another interesting insight from our study is the structured C-terminus of β-tubulin. The C-termini of tubulin in the doublet normally have polyglycylation and polyglutamylation, in particular, the B-tubule (47). In reconstituted microtubules and other places in the doublet, the C-termini are highly flexible and cannot be visualized. However, we observed the C-terminus of β-tubulin in PF A1 which appears to interact with PACRG and FAP20. In addition, the position of FAP126 and FAP106 binding on top of tubulin molecules also suggest they are interacting with the C-termini of tubulins. In vitro study shows that the C-tails of tubulins must be suppressed for the outer junction to be formed (14). This suggests that the C-terminus might have a role in the assembly and or stability of the doublet. Defects in tubulin polyglutamylase enzyme have indeed led to partially formed B-tubules (48). This could indicate a role for polyglutamylation in the interaction and recruitment at the IJ PF, specifically PACRG and FAP20. Lack of polyglutamylation can lead to an easily detachable PACRG and FAP20 and hence the partial assembly of the B-tubule. Finally, it is possible that MIPs can act as readers for the post-translational modifications of tubulin for their orderly recruitment and assembly.

## MATERIALS AND METHODS

### Preparation of doublet samples

WT Chlamydomonas cells (cc124) were obtained from *Chlamydomonas* source center and cultured either on Tris-acetatephosphate (TAP) media with shaking or stirring with 12 hr light-12 hr dark cycle. For flagella purification, *Chlamydomonas* cells were cultured in 1.5 L of liquid TAP media with stirring until OD600 reached around 0.5-0.6 and harvested by low-speed centrifugation (700g for 7 min at 4°C). *Chlamydomonas* flagella were purified by dibucaine method (49), resuspended in HMDEKP buffer (30 mM HEPES, pH 7.4, 5 mM MgSO4, 1 mM DTT, 0.5 mMc, 25 mM Potassium Acetate, 0.5% polyethylene glycol, MW 20,000) containing 10 μM paclitaxel, 1 mM PMSF, 10 μg/ml aprotinin and 5 μg/ml leupeptin. Paclitaxel was added to the buffer since *Chlamydomonas* doublets were more vulnerable to high salt extraction compared with *Tetrahymena* doublets (data not shown). Isolated flagella were demembraned by incubating with HMDEKP buffer containing final 1.5% NP40 for 30 min on ice. After NP40 treatment, *Chlamydomonas* doublets were incubated with final 1 mM ADP for 10 min at room temperature to activate dynein and then incubated with 0.1 mM ATP for 10 min at room temperature to induce doublet sliding. Since the *Chlamydomonas* doublets were harder to split compared to *Tetrahymena* doublet, sonication was done before ADP/ATP treatment. After this, *Chlamydomonas* doublets were incubated twice with HMDEKP buffer containing 0.6 M NaCl for 30 min on ice, spinned down (16,000 g and 10 minutes), and resuspended. *Chlamydomonas* doublets were not dialyzed against low salt buffer since it was difficult to remove radial spokes. The gel of purification process is in Supplementary Fig. 1A.

Tetrahymena doublets were isolated according to our previous work (7, 8).

### Cryo-electron microscopy

3.5 ul of sample of doublets (∼4 mg/ml) was applied to a glow-discharged holey carbon grid (Quantifoil R2/2), blotted and plunged into liquid ethane using Vitrobot Mark IV (Thermo Fisher Scientific) at 25°C and 100% humidity with a blot force 3 or 4 and a blot time of 5 sec. 9,528 movies were obtained on a Titan Krios (Thermo Fisher Scientific) equipped with Falcon II camera at 59,000 nominal magnification. The pixel size was 1.375 Å/pixel. Dataset for *Tetrahymena* was described in Ichikawa et al. (8). *Chlamydomonas* dataset was collected with a dose of 28-45 electron/Å2 with 7 frames. The defocus range was set to between -1.2 and -3.8 um.

The Chlamydomonas doublet structures were performed according to Ichikawa et al. (8). In short, movies were motion corrected using MotionCor2 (50). The contrast transfer function were estimated Gctf (51). The doublets were picked using e2helixboxer (52).

270,713 and 122,997 particles were used for the 16-nm reconstruction of 48-nm repeating unit of *Chlamydomonas*. 279,850 particles were used for the 16-nm reconstruction of *Tetrahymena*. The final Gold Standard FSC resolutions of the 16-nm and 48-nm reconstruction for Chlamydomonas after contrast transfer function refinement and polishing using 0.143 FSC criterion in Relion3 (53) are 4.5 and 3.8 Å, respectively. Using focus refinement of the IJ of the 16-nm reconstruction for Chlamydomonas, the resolution reaches 3.6 Å resolution. The resolution for the 16-nm reconstruction of Tetrahymena was 3.6 Å. Focus refinement of the IJ of Tetrahymena did not improve the resolution of the IJ due to the flexibility of this region. The maps were local sharpened (8). Local resolution estimation was performed using MonoRes (54).

### Modelling

#### C. reinhardtii α-β-tubulin

A homology model of *C. reinhardtii* α-β-tubulin (Uniprot sequence α: P09204, β: P04690) was constructed in Modeller v9.19 (55) using PDB 5SYF as template. The model was refined using real-space refinement (56) and validated using comprehensive validation for cryo-EM in Phenix v1.16 (57).

#### PACRG and FAP20

A partial homology model of *C. reinhardtii* PACRG (B1B601) was constructed using the crystal structure of the human homolog (Q96M98-1) as template (58). The model was completed by building segments N2-D148 and Y249-L270 de novo in density using Coot v0.8.9.1 (59). The density for segment M89-K101 is missing, likely due to flexibility in this region. C. reinhardtii FAP20 (A8IU92) was completely built de novo in density. Both models were refined and validated as described for α-β-tubulin.

#### FAP52

The density was traced in Coot v0.8.9.1 (59) according to a double beta-propeller topology similar to PDB 2YMU, which agrees with the I-TASSER (60) tertiary structure prediction of FAP52 (Uniprot: A0A2K3D260). The bulky residues of FAP52 were used as anchors to maintain the correct registry in lower resolution areas. The model has been overfit on the left side (segment D341-P627) where the density signal is significantly lower, likely due to heterogeneity. The final model was refined and validated as described above.

#### FAP276

The density for FAP276 was segmented and traced to around 80 amino acids and ∼9 kDa in mass. Candidates from the wild type mass spectrometry data were compared to the FAP52 knockout data and reduced to only FAP276, which was completely missing in the latter. The secondary structure prediction (61) as well as the sequence of FAP276 (Phytozome: Cre04.g216250) agree unambiguously with the density signature in that region. The model was traced, refined and validated as described above.

#### FAP126

The density for the hook, which is mostly disordered, was traced as before to 133 amino acids and ∼15 kDa. The density had clear side chains signature, particularly in an area where it appeared to have a Trp residue followed by a Pro, four more amino acids and another Trp. Doing a local search against the entire Chlamydomonas reinhardtii proteome (Uniprot: UP000006906), in both C- and N-termini directions, using a regular expression matching the pattern above, gives a single hit: FAP126. Furthermore, inspecting candidates in the wild type mass spectrometry that has a similar abundance to IJ proteins after normalizing the quantitative peptide value by the molecular weight places FAP126 in the top list of candidates for this density. As before, the sequence has matching secondary structure prediction and unambiguous density signature agreement throughout the entire sequence. The model was modelled and refined as mentioned above.

### Inter-PF angle (lateral curvature) measurement

The inter-PF angle between each PF pair are measured according to Ichikawa et al., (7).

### Visualization

The maps and models were segmented, coloured and visualized using Chimera (62) and ChimeraX (63).

### Mass spectrometry

Sample preparation and mass spectrometry of FAP52 mutant and relative quantification compared to wild type *Chlamydomonas* was done according to Dai et al., 2019. (25). The ratio between the averaged quantitative values from the mass spectrometry (n=3) and a proteins molecular weight was used to calculate their stoichiometry in the axoneme.

### Transcriptomics analysis

Transcriptomics analysis of PACRG, FAP20, FAP52, FAP126, FAP106, FAP45 and DCX using consensus normalized expression levels for 55 tissue types and 7 blood cell types was done according to (29).

## DATA AVAILABILITY

Cryo-EM maps have been deposited in EM data bank (EMDB) with accession numbers of EMD-XXXXX (48-nm averaged Chlamydomonas doublet), EMD-XXXXX (16-nm averaged Chlamydomonas IJ region) and EMD-XXXXX (16-nm averaged Tetrahymena IJ region). The model of IJ is available in Protein Data Bank (PDB) with an accession number of PDB: XXXX.

## ACKNOWLEDGEMENT

We acknowledge the Facility for Electron Microscopy Research of McGill University where our cryo-EM experiments were conducted, particularly, Drs. Kaustuv Basu and Kelly Sears. We are indebted to Dr. Masahide Kikkawa for sharing the FAP52 knockout Chlamydomonas strain. This research was financially supported by Natural Sciences and Engineering Research Council of Canada (RGPIN-2016-04954), Canada Institute of Health Research (CIHR PJT-156354) and the Canada Institute for Advanced Research Arzieli Global Scholars Program to K.H.B., M.I. and S.K. were supported by Japan Society for the Promotion of Science (JSPS) for JSPS Overseas Research Fellowships and JSPS Overseas Challenge Program for Young Researchers, respectively. A.A.Z.K. is supported by CIHR and the Al Ghurair Foundation for Education. We declare no competing financial interests.

## Notes

#### Summary of Updates

New results regarding FAP126 and FAP106. Therefore, one more figure is added and the result & discussion are changed significantly.

