## Supplementary Materials for "The inner junction complex of the cilia is an interaction hub that involves tubulin post-translational modifications"

**Figure S1:** Data related to the doublet structures

(A) Schematics of fractionation of the axoneme in this study. Doublets were split from axoneme, and outside proteins were removed to obtain a simpler sample for cryo-EM. (B) A typical cryo-EM image of *Chlamydomonas* doublets. (C) Gold-standard Fourier Shell Correlation of the 48-nm repeat and 16-nm repeat doublet maps of *Chlamydomonas* and *Tetrahymena*. (D) Local resolution estimation of the 16-nm repeat maps from *Chlamydomonas* and *Tetrahymena* using MonoRes. The resolution of the B-tubule in *Tetrahymena* is lower due to the loss of the IJ PF. (E) The remaining PACRG- and FAP20-like densities in the *Tetrahymena* doublet structure. (F) Superimposition of the tomographic structure of the intact doublet (EMD-2132) with the 48-nm structure of the *Chlamydomonas* doublet in this study. The DRC is colored green. (G) Enlarged view of the missing PACRG unit at the IJ PF, where the DRC binds.

**Figure S2:** Atomic models of PACRG, FAP20, FAP52 and FAP276.

Atomic model of (A) PACRG, (C) FAP20, (E) FAP52 and (G) FAP276. Illustration of the cryo-EM density quality at selected regions of (B) PACRG, (D) FAP20, (F) FAP52 and (H) FAP276.

**Figure S3:** Multiple sequence alignment of FAP20 shows that it is highly conserved.

**Figure S4:** Data related to FAP52.

(A) Atomic model of *Chlamydomonas* FAP52 from inside the *Chlamydomonas* density map. (B) Atomic model of *Chlamydomonas* FAP52 fitted inside the *Tetrahymena* map highlights the longer loop (red) from *Chlamydomonas*. (C) Alignment of FAP52 from several species shows that *Chlamydomonas* has a longer loop in one beta propeller blade. The long loop is responsible for the interaction with PACRG. (D, E) α-K40 loop from PF B9 in *Chlamydomonas*. (F, G) Superimposition of the acetylated α-K40 loops from *Chlamydomonas* PF B9, B10 and *Tetrahymena* A12.

**Figure S5:** Data related to the Tether densities.

(A-B) The small helix (indicated by the dashed box) from *Chlamydomonas* (A) and *Tetrahymena* Tether loop appears to interact with α-tubulin. (C) Multiple sequence alignment of FAP106, the candidate for the Tether loop from a few organisms. (D) Secondary structure prediction of FAP106. The big cylinder represents helical prediction. Some of the beta-sheets are omitted since it is not easy to match beta-sheet with densities.

**Figure S6:** Data related to FAP126.

(A) Multiple sequence alignment of FAP126 from a few organisms. FAP126 of *Chlamydomonas* still have the SH3 binding domain while lacks the proline-rich region compared to other species. (B) Atomic model of FAP126 fitted inside its segmented density. (C) Table of pairwise correlation coefficients between tissue mRNA expression levels, color-coded from low (red) to high (blue) values. ENKUR is the homolog of FAP106 in human. DCX is a microtubule associated protein in neuron, picked as a control. (D) Correlation graphs of consensus normalized expression levels for two selected pairs of genes (PACRG/FAP20 and PACRG/FAP52). Tissues showing high levels of expression of one or both genes are labeled. Correlation coefficients (*r*) are indicated.

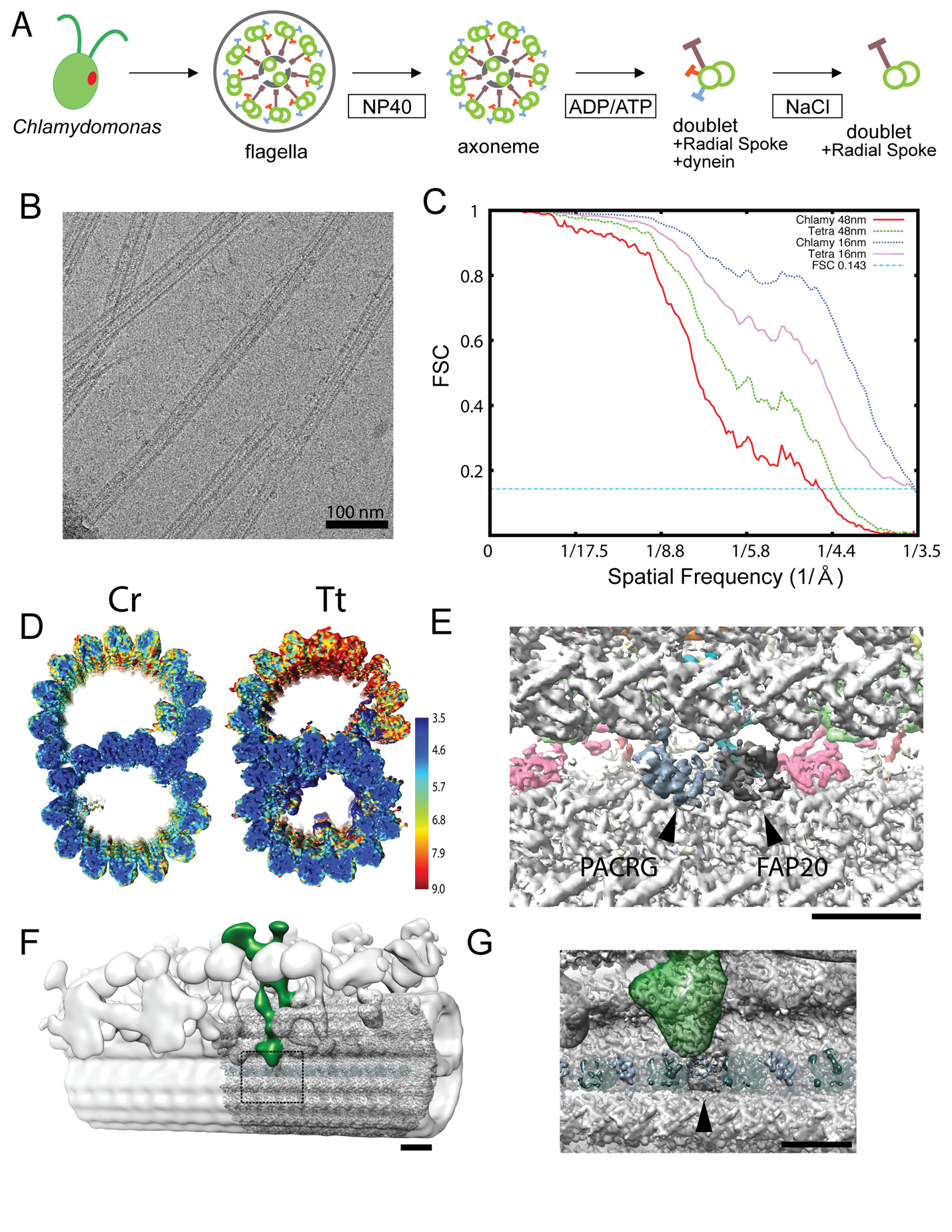

**Figure S1:** Data related to the doublet structures

(A) Schematics of fractionation of the axoneme in this study. Doublets were split from axoneme, and outside proteins were removed to obtain a simpler sample for cryo-EM. (B) A typical cryo-EM image of *Chlamydomonas* doublets. (C) Gold-standard Fourier Shell Correlation of the 48-nm repeat and 16-nm repeat doublet maps of *Chlamydomonas* and *Tetrahymena*. (D) Local resolution estimation of the 16-nm repeat maps from *Chlamydomonas* and *Tetrahymena* using MonoRes. The resolution of the B-tubule in *Tetrahymena* is lower due to the loss of the IJ PF. (E) The remaining PACRG- and FAP20-like densities in the *Tetrahymena* doublet structure. (F) Superimposition of the tomographic structure of the intact doublet (EMD-2132) with the 48-nm structure of the *Chlamydomonas* doublet in this study. The DRC is colored green. (G) Enlarged view of the missing PACRG unit at the IJ PF, where the DRC binds.

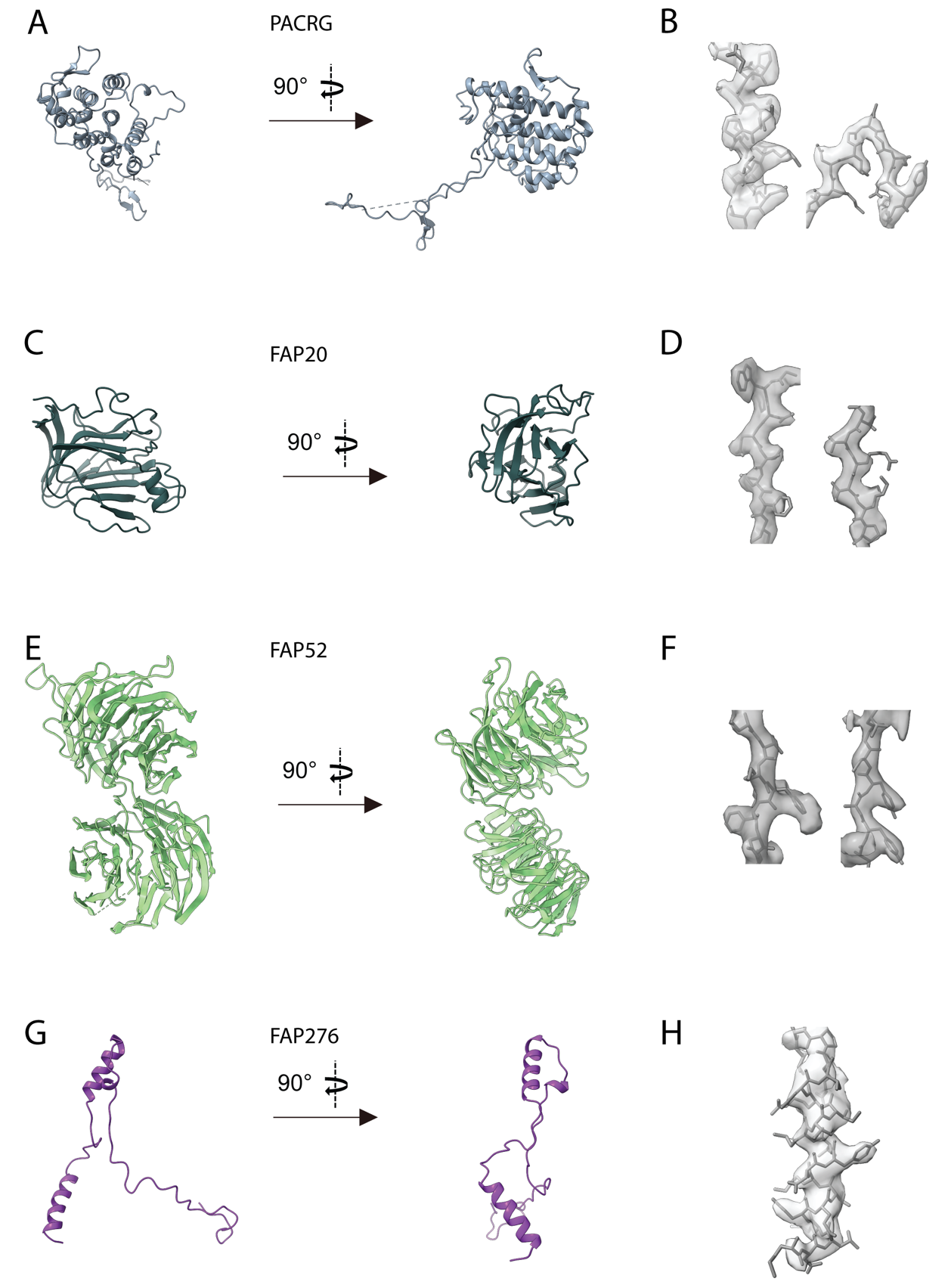

**Figure S2:** Atomic models of PACRG, FAP20, FAP52 and FAP276.

Atomic model of (A) PACRG, (C) FAP20, (E) FAP52 and (G) FAP276. Illustration of the cryo-EM density quality at selected regions of (B) PACRG, (D) FAP20, (F) FAP52 and (H) FAP276.

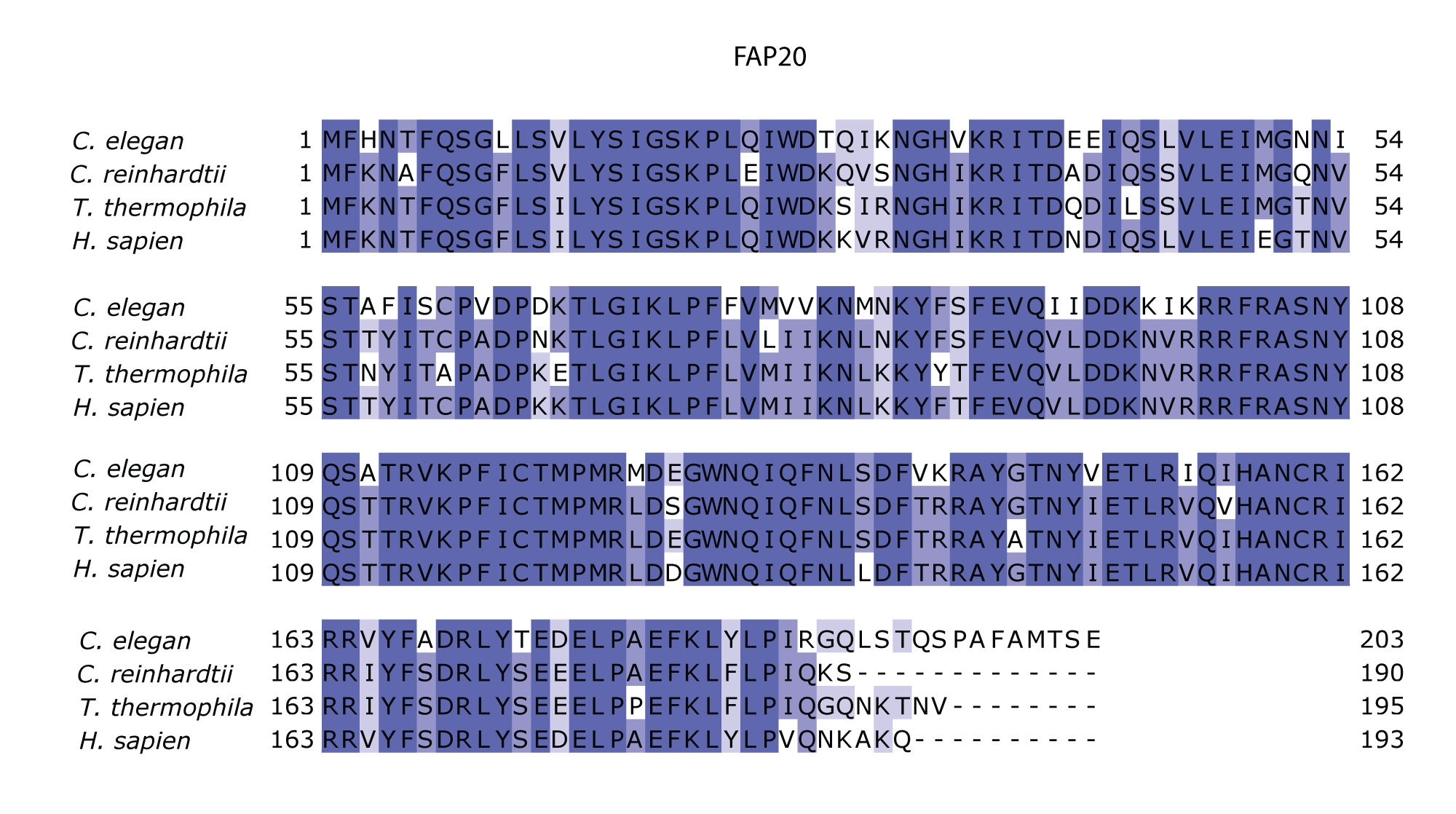

**Figure S3:** Multiple sequence alignment of FAP20 shows that it is highly conserved.

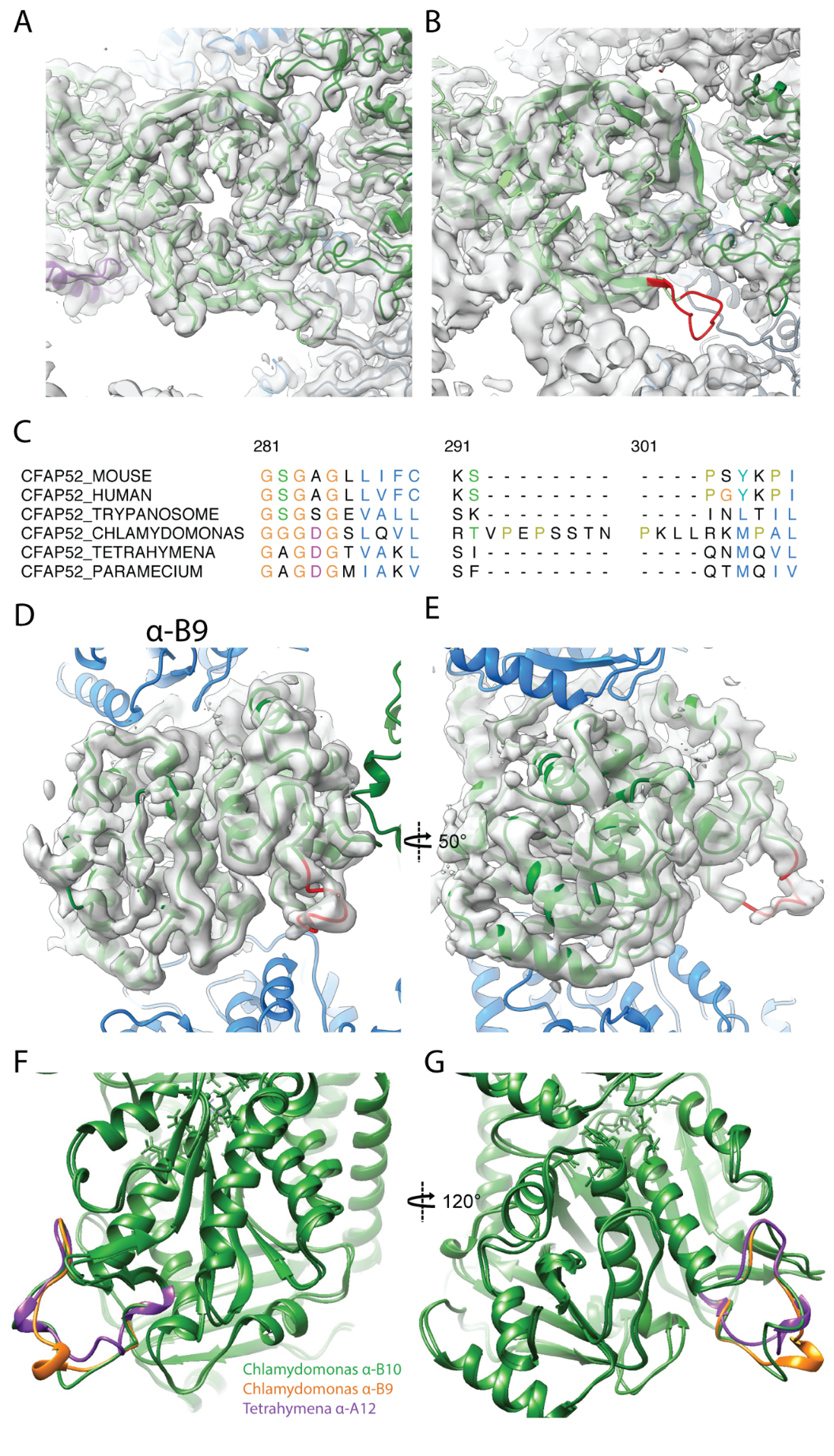

**Figure S4:** Data related to FAP52.

(A) Atomic model of *Chlamydomonas* FAP52 from inside the *Chlamydomonas* density map. (B) Atomic model of *Chlamydomonas* FAP52 fitted inside the *Tetrahymena* map highlights the longer loop (red) from *Chlamydomonas*. (C) Alignment of FAP52 from several species shows that *Chlamydomonas* has a longer loop in one beta propeller blade. The long loop is responsible for the interaction with PACRG. (D, E) α-K40 loop from PF B9 in *Chlamydomonas*. (F, G) Superimposition of the acetylated α-K40 loops from *Chlamydomonas* PF B9, B10 and *Tetrahymena* A12.

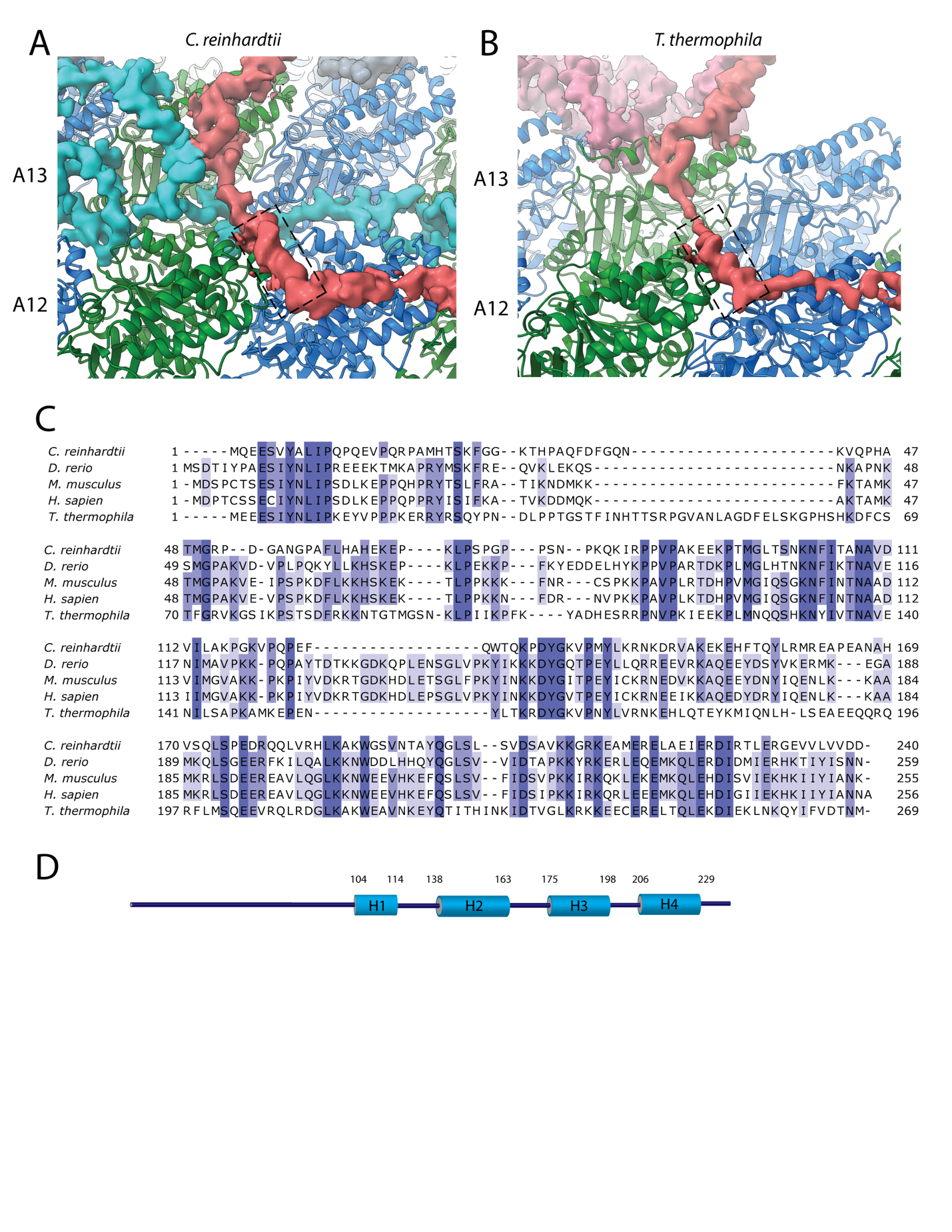
**Figure S5:** Data related to the Tether densities.

(A-B) The small helix (indicated by the dashed box) from *Chlamydomonas* (A) and *Tetrahymena* Tether loop appears to interact with α-tubulin. (C) Multiple sequence alignment of FAP106, the candidate for the Tether loop from a few organisms. (D) Secondary structure prediction of FAP106. The big cylinder represents helical prediction. Some of the beta-sheets are omitted since it is not easy to match beta-sheet with densities.

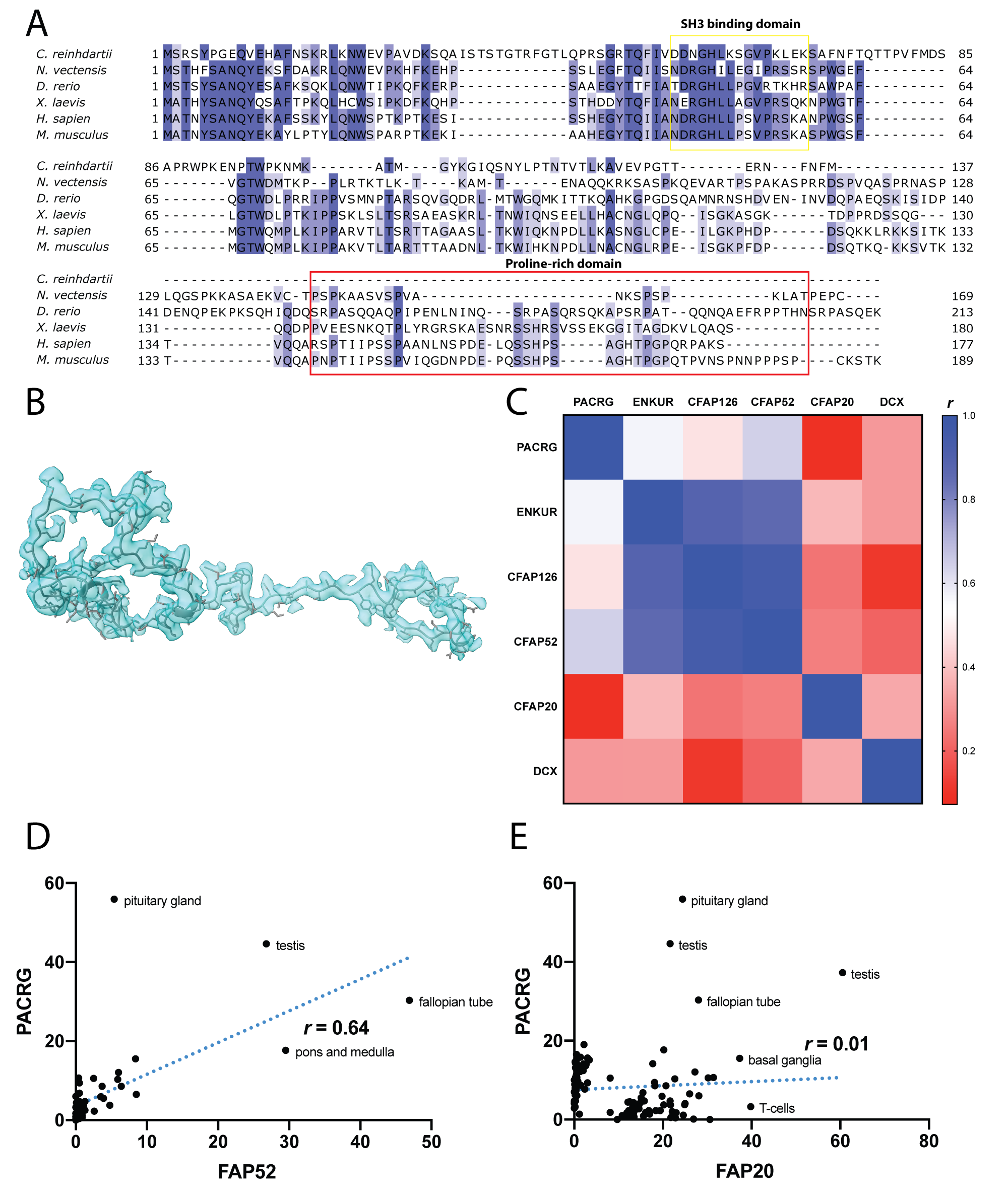

**Figure S6:** Data related to FAP126.

(A) Multiple sequence alignment of FAP126 from a few organisms. FAP126 of *Chlamydomonas* still have the SH3 binding domain while lacks the proline-rich region compared to other species. (B) Atomic model of FAP126 fitted inside its segmented density. (C) Table of pairwise correlation coefficients between tissue mRNA expression levels, color-coded from low (red) to high (blue) values. ENKUR is the homolog of FAP106 in human. DCX is a microtubule associated protein in neuron, picked as a control. (D) Correlation graphs of consensus normalized expression levels for two selected pairs of genes (PACRG/FAP20 and PACRG/FAP52). Tissues showing high levels of expression of one or both genes are labeled. Correlation coefficients (*r*) are indicated.

**SUPPLEMENTARY TABLE**

**Supplementary Table 1:** Significantly reduced or missing proteins in FAP52 compared to WT using relative mass spectrometry quantification.

| Names | Uniprot ID | **WT**  exclusive unique peptide counts (quantitative values after normalization) | **FAP52**  exclusive unique peptide counts (quantitative values after normalization) | **FAP52/WT ratio**  (quantitative values were used) | **p-values**  **(WT vs FAP52)** | **Log2(Fold Change *(*FAP52/WT))** |
| --- | --- | --- | --- | --- | --- | --- |
| **FAP20** | **A8IU92** | **14, 10, 12 (27, 19, 33)** | **14, 12, 13 (33, 17, 18)** | **0.86** | **0.65** | **-0.31** |
| **FAP45** | **A8I9E8** | **31, 27, 12 (60, 37, 33)** | **31, 30, 30 (48, 43, 40)** | **1.01** | **0.96** | **0.016** |
| **PACRG** | **A8I2Z6** | **13, 9, 10 (41, 26, 48)** | **15, 13, 13 (70, 42, 38)** | **1.30** | **0.38** | **0.38** |
| **Tektin** | **A8J8F6** | **20, 22, 14 (60, 61, 52)** | **29, 24 24 (74, 45, 43)** | **0.88** | **0.74** | **-0.096** |
| ARL3 | A8ISN6 | 2, 2, 1 (2, 1, 2) | 0, 0, 0 (0, 0, 0) | 0.0 | 0.0013 | -10.0 |
| CHLREDRAFT_171815 | A8HQQ4 | 2, 6, 1 (2, 5, 2) | 0, 0, 0 (0, 0, 0) | 0.0 | 0.035 | -10.0 |
| CHLREDRAFT_156073 | A8I1U2 | 1, 1, 1 (2, 1, 2) | 0, 0, 0 (0, 0, 0) | 0.0 | 0.024 | -10.0 |
| FAP276 | A8J9P2 | 3, 3, 2 (8 ,7, 14) | 0, 0, 0 (0, 0, 0) | 0.0 | 0.015 | -10.0 |
| CFAP52 | A8ILK1 | 27, 21, 15 (59, 73, 105) | 0, 0, 0 (0, 0, 0) | 0.0 | 0.0046 | -10.0 |
| FAP36 | A8IZX7 | 3, 3, 1 (3, 3, 2) | 0, 0, 0 (0, 0, 0) | 0.0 | 0.0023 | -10.0 |
| CrCDPK1 | A8IHF4 | 3, 5, 2 (3, 4, 4) | 0, 0, 0 (0, 0, 0) | 0.0 | <0.0001 | -10.0 |
| CHLREDRAFT_176830 | A8J922 | 2, 1, 2 (2, 1, 1) | 0, 0, 0 (0, 0, 0) | 0.0 | 0.024 | -10.0 |
| FAP173 | A8JAF7 | 3, 3, 1 (5, 3, 2) | 0, 0, 0 (0, 0, 0) | 0.0 | 0.012 | -10.0 |
| FAP29 | A8J3X6 | 2, 3, 2 (3, 3, 4) | 0, 0, 0 (0, 0, 0) | 0.0 | 0.00045 | -10.0 |
| CHLREDRAFT_181390 | A8JJY2 | 1, 1, 1 (1, 1, 2) | 0, 0, 0 (0, 0, 0) | 0.0 | 0.028 | -10.0 |
| ANK2 | A8HNK2 | 1, 2, 1 (1, 1, 2) | 0, 0, 0 (0, 0, 0) | 0.0 | 0,0041 | -10.0 |
| FAP5 | A8JAI0 | 11, 15, 7 (23, 22, 23) | 2, 0, 0 (1, 0, 0) | 0.014 | <0.0001 | -5.5 |
| FAP164 | A8JC79 | 4, 5, 1 (6, 4, 2) | 0, 0, 1 (0, 0, 0) | 0.0 | 0.022 | -5.1 |
| FAP288 | A8IJV3 | 13, 12, 9 (18, 14, 23) | 2, 1, 1 (1, 0, 0) | 0.018 | 0.0021 | -4.6 |
| CHLREDRAFT_177061 | A8J9A4 | 7, 7, 2 (8, 6, 4) | 0, 1, 1 (0, 0, 0) | 0.0 | 0.0071 | -4.5 |
| TEF20 | A8IL00 | 2, 1, 1 (2, 1, 2) | 0, 1, 0 (0, 0, 0) | 0.0 | 0.036 | -3.6 |
| CHLREDRAFT_191579 | A8J3S1 | 2, 4, 2 (3, 4, 4) | 1, 0 ,1 (1, 0 , 0) | 0.09 | 0.0003 | -3.4 |
| CHLREDRAFT_111330 | A8IAY6 | 2, 2, 1 (3, 1, 2) | 1, 0, 0 (1, 0, 0) | 0.16 | 0.03 | -3.2 |
| 14-3-3 | Q7X7A7 | 9, 9, 3 (15, 10, 6) | 5, 0, 1 (4, 0, 0) | 0.13 | 0.03 | -3.0 |
| Isocitrate lyase | A8J244 | 12, 8, 9 (23, 8, 19) | 6, 2, 2 (5, 1, 1) | 0.14 | 0.036 | -2.9 |
| CHLREDRAFT_141580 | A8I9N1 | 7, 11, 4 (13, 16, 17) | 3, 3, 3 (3, 2, 2) | 0.19 | 0.0006 | -2.7 |
| Elongation Factor 2 | A8JHX9 | 15,17,5 (27, 22, 10) | 3, 6, 6 (3, 3, 3) | 0.15 | 0.026 | -2.7 |
| CHLREDRAFT_189452 | A8IT59 | 4, 6, 1 (5, 5, 2) | 1, 1, 1 (1, 0, 1) | 0.17 | 0.026 | -2.7 |
| CHLREDRAFT_175290 | A8J364 | 4, 3, 2 (5, 3, 6) | 1, 2, 2 (1, 1, 1) | 0.21 | 0.014 | -2.6 |
| CHLREDRAFT_111269 | A8IBY2 | 1, 4, 1 (1, 3, 2) | 0, 2, 1 (0, 1, 0) | 0.17 | 0.045 | -2.4 |
| FAP138 | A8IUQ2 | 4, 6, 3 (8, 7, 6) | 2, 3, 3 (1, 1, 1) | 0.14 | 0.00046 | -2.4 |
| FAP31 | A8JDM7 | 6, 8, 4 (8, 7, 8) | 7, 0, 0 (5, 0, 0) | 0.21 | 0.024 | -2.2 |
| IFT80 | A8IXE2 | 4, 5, 1 (5, 4, 2) | 1, 1, 2 (1, 1, 1) | 0.27 | 0.022 | -2.2 |
| CHLREDRAFT_189792 | A8HPX1 | 16, 21, 6 (18, 17, 12) | 0, 11, 13 (0, 6, 5) | 0.23 | 0.0096 | -2.1 |
| CHLREDRAFT_206178 | A8IP72 | 17,  24, 10 (20, 19,23) | 0, 15, 18 (0, 7, 8) | 0.24 | 0.0053 | -2.0 |
| GSK3 | Q6IV67 | 4, 3, 2 (5, 2, 4) | 2, 2, 1 (1, 1, 0) | 0.18 | 0.030 | -1.8 |
| CHLREDRAFT_144025 | A8IF86 | 13, 27, 14 (16, 27, 35) | 14, 12, 13 (11, 7, 7) | 0.32 | 0.035 | -1.7 |
| FAP148 | A8IAT9 | 29, 41, 11 (44, 52, 27) | 14, 22, 26 (14, 12, 15) | 0.33 | 0.022 | -1.6 |
| FAP85 | A8J250 | 7, 11, 6 (14, 15, 19) | 8, 4, 4 (10, 3, 2) | 0.31 | 0.024 | -1.6 |
| Phototropin | A8IXU7 | 30, 22, 18 (56, 44, 64) | 25, 14, 17 (41,12, 13) | 0.40 | 0.041 | -1.3 |
| p38 | A4PET3 | 3, 8, 4 (7 ,7, 8) | 5, 3, 3 (5, 2, 1) | 0.36 | 0.019 | -1.3 |
| FAP39 | A8J0V2 | 6, 10, 4 (13, 11, 8) | 9, 5, 4 (7, 4, 3) | 0.42 | 0.022 | -1.1 |

**Supplementary Movie 1:** The IJ complex of the *Chlamydomonas reinhardtii* flagella.
